# PLEKHA5, PLEKHA6 and PLEKHA7 bind to PDZD11 to target the Menkes ATPase ATP7A to the cell periphery and regulate copper homeostasis

**DOI:** 10.1101/2021.06.17.448791

**Authors:** Sophie Sluysmans, Isabelle Méan, Tong Xiao, Amina Boukhatemi, Flavio Ferreira, Lionel Jond, Christopher J. Chang, Sandra Citi

## Abstract

Copper homeostasis is crucial for cellular physiology and development, and its dysregulation leads to disease. The Menkes ATPase ATP7A plays a key role in copper efflux, by trafficking from the Golgi to the plasma membrane upon cell exposure to elevated copper, but the mechanisms that target ATP7A to the cell periphery are poorly understood. PDZD11 interacts with the C-terminus of ATP7A, which contains sequences involved in ATP7A trafficking, but the role of PDZD11 in ATP7A localization is unknown. Here we identify PLEKHA5 and PLEKHA6 as new interactors of PDZD11, which similarly to the junctional protein PLEKHA7 bind to PDZD11 N-terminus through their WW domains. Using CRISPR-KO kidney epithelial cells, we show by immunofluorescence that WW-PLEKHAs (PLEKHA5, PLEKHA6, PLEKHA7) recruit PDZD11 to distinct plasma membrane localizations, and that they are required for the efficient anterograde targeting of ATP7A to the cell periphery in elevated copper. Pulldown experiments show that WW-PLEKHAs promote PDZD11 interaction with the C-terminus of ATP7A. However, WW-PLEKHAs and PDZD11 are not necessary for ATP7A Golgi localization in basal copper, ATP7A copper-induced exit from the Golgi, and ATP7A retrograde trafficking to the Golgi. Finally, measuring bioavailable copper with the labile copper probe CF4 shows that WW-PLEKHAs and PDZD11 are required to maintain low intracellular copper levels when cells are exposed to elevated copper. These data indicate that WW-PLEKHAs-PDZD11 complexes regulate the localization and function of ATP7A to modulate cellular copper homeostasis.

## Introduction

Copper is an essential micronutrient as a cofactor of vital oxidative enzymes, and both its deficiency and overload are pathological, as exemplified by the Menkes and Wilson diseases (Bandmann *et al*, 2015; Czlonkowska *et al*, 2018). The brain and nervous tissue have a particularly high oxidative demand and requirement for copper-dependent enzyme activities, and several neurological disorders, including Alzheimer, Parkinson and Huntingdon diseases, are associated with disrupted copper homeostasis (Ackerman & Chang, 2018; Kaler, 2011; Lutsenko *et al*, 2019; Telianidis *et al*, 2013; Zlatic *et al*, 2015). Intracellular copper is physiologically finely tuned through the coordination of copper uptake, intracellular trafficking by copper chaperones, and efflux by specific copper efflux transporters (reviewed in (Camakaris *et al*, 1999; Hartwig *et al*, 2019; La Fontaine & Mercer, 2007; Lutsenko *et al*, 2007;

Polishchuk & Lutsenko, 2013)). The P-type ATPases ATP7A (Menkes copper ATPase) and ATP7B (Wilson copper ATPase) play a major role in copper efflux in different tissues (La Fontaine & Mercer, 2007; Lutsenko *et al*., 2007; Nevitt *et al*, 2012). Under basal copper conditions ATP7A is localized in a specific compartment of the Trans-Golgi Network (TGN), whereas elevated copper levels induce its translocation from the TGN to a compartment of post-Golgi membrane vesicles which are trafficked to the plasma membrane and whose exocytosis and endocytic recycling drive copper efflux (Camakaris *et al*, 1995; Cobbold *et al*, 2002; Dierick *et al*, 1997; Holloway *et al*, 2007; La Fontaine *et al*, 1999; La Fontaine *et al*, 1998; Monty *et al*, 2005; Nyasae *et al*, 2007; Petris & Mercer, 1999; Petris *et al*, 1996; Petris *et al*, 2002; Yamaguchi *et al*, 1996).

Although the severity of Menkes disease correlates with the intracellular localization of ATP7A (Skjorringe *et al*, 2017), the molecular mechanisms that regulate the trafficking and plasma membrane localization of ATP7A-containing vesicles remain unresolved. In cultured polarized epithelial cells C-terminal di-leucine and PDZ-binding motifs are required for basolateral plasma membrane targeting of ATP7A in elevated copper (Greenough *et al*, 2004). The single PDZ domain-containing protein (PDZD11, also known as AIPP1) was identified as an interactor of the C-terminal 15 amino acids of ATP7A and in *Saccharomyces cerevisiae* this interaction was not dependent on copper levels (Stephenson *et al*, 2005). Although these observations suggest that PDZD11 could be involved in plasma membrane targeting of ATP7A, the role of PDZD11 in ATP7A trafficking and in copper homeostasis has not been explored.

We identified PDZD11 as an interactor of the adherens junction (AJ) protein PLEKHA7 (Guerrera *et al*, 2016). The N-terminal tandem WW domains of PLEKHA7 bind to the N-terminal proline-rich sequence of PDZD11 to recruit PDZD11 to AJ, and this interaction is required for the accumulation of the transmembrane proteins nectins, tetraspanin-33 (Tspan33) and ADAM10 at AJ (Guerrera *et al*., 2016; Rouaud *et al*, 2020; Shah *et al*, 2018). Although we detected PDZD11 labeling predominantly at cell-cell junctions using antibodies against the endogenous protein (Guerrera *et al*., 2016), PDZD11 was reported to interact not only with ATP7A (Stephenson *et al*., 2005), but also with additional transmembrane transporters, such as the plasma membrane calcium ATPase (PMCA) and the sodium-dependent multivitamin transporter SLC5A6 (Goellner *et al*, 2003; Nabokina *et al*, 2011), which are not localized at junctions, but along the basolateral membrane of polarized epithelial cells (Chicka & Strehler, 2003;

Greenough *et al*., 2004; Subramanian *et al*, 2009). This suggests that PDZD11 interacts with these other ligands at these sites independently of PLEKHA7. To address this question and gain more insight into the molecular complexes that implicate PDZD11, we searched for new PDZD11 interactors through a Yeast-2-hybrid (Y2H) screen. We report here that PLEKHA5 (Pleckstrin homology domain-containing family A member 5) and PLEKHA6 (Pleckstrin homology domain-containing family A member 6), also known respectively as phosphatidylinositol-three-phosphate-binding PH-domain protein-2 (PEPP2) and -3 (PEPP3) (Dowler *et al*, 2000), are new PDZD11- interacting proteins, which are localized in the cytoplasm and near the apical, lateral and basal plasma membranes in epithelial cells within tissues and in culture. Since PDZD11 binds to ATP7A sequences required for its trafficking to the cell periphery (Greenough *et al*., 2004; Stephenson *et al*., 2005), we hypothesized that PDZD11, PLEKHA5, PLEKHA6 and PLEKHA7 are involved in ATP7A trafficking. Here we validate this hypothesis, and we show that PLEKHA5, PLEKHA6 and PLEKHA7 promote the binding of PDZD11 to the C-terminus of ATP7A and together with PDZD11 are also required for efficient copper efflux upon cell exposure to elevated copper.

## Results

### PLEKHA5 and PLEKHA6 are new interactors of PDZD11

In addition to interactions with known partners, e.g. PLEKHA7 (residues 1-113 of transcript variant NM_175058.4) (Guerrera *et al*., 2016) and the sodium-dependent multivitamin transporter SLC5A6 (Nabokina *et al*., 2011), a Y2H screen using full-length PDZD11 as a bait revealed high confidence interactions with the N-terminal sequences of PLEKHA5 (residues 1-117 of transcript variant 1, NM_019012.5) and PLEKHA6 (residues 1-93 of transcript variant X11, XM_011509297.2) (Fig. 1A). Sequence analysis shows that similarly to PLEKHA7, and unlike other members of the PLEKHA family of proteins, these PLEKHA5 and PLEKHA6 isoforms contain N-terminal tandem WW domains. The highest degree of sequence similarity is displayed by the N-terminal regions, comprising the WW and PH domains (Fig. S1A-E). We propose the name WW-PLEKHAs for these members of the PLEKHA family, which contain tandem WW domains. WW-PLEKHAs also comprise a PH domain, typical of the PLEKHA family, and C-terminal regions that contain coiled-coil (CC) and proline-rich (Pro-rich) domains (Fig. 1A, Fig. S1A).

**Figure 1.**
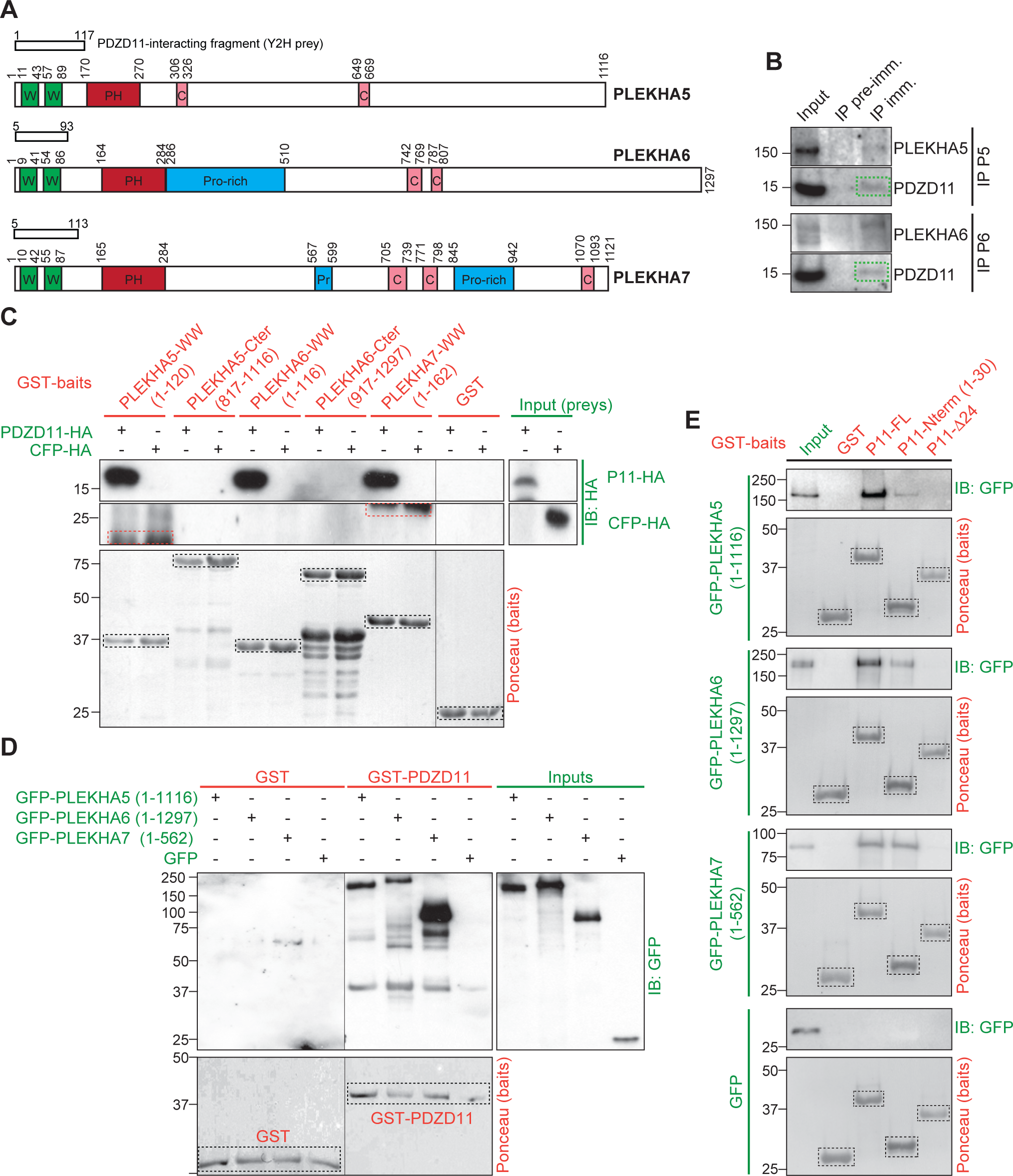
PLEKHA5 and PLEKHA6 interact with the N-terminus of PDZD11 through their tandem WW domains. (A) Schemes of human PLEKHA5, PLEKHA6 and PLEKHA7 showing amino acid positions and structural domains: WW (Trp-Trp) green, PH (pleckstrin homology) red, Proline-rich (Pro-rich) blue, coiled-coil (CC) pink. Boxes show yeast-2-hyrid (Y2H) screen preys. (B) IB analysis of PDZD11 in immunoprecipitates (IP) of either PLEKHA5 (P5) or PLEKHA6 (P6) from lysates of Caco-2 cells (pre-immune serum as negative control). (C) IB analysis, using anti-HA antibody, of GST pull-downs of either PDZD11-HA (P11-HA) or CFP-HA (negative control) (preys, green), using as baits (red) either GST or GST fused to the indicated sequences. Bait degradation products are non-specifically labeled in the CFP-HA pulldown (red dashed rectangles). (D) IB analysis, using anti-GFP antibodies, of GST pull-downs of GFP-tagged PLEKHA5 (full-length), PLEKHA6 (full-length) and PLEKHA7 (N-ter.) preys using as baits either GST or GST-PDZD11. (E) IB analysis, using anti-GFP antibodies, of GST pull-downs of GFP-tagged PLEKHA5 (full-length), PLEKHA6 (full-length) and PLEKHA7 (N-ter.) preys (green) using as baits (red) either GST, or GST-PDZD11 full-length, or the N-terminal 30 residues (P11-Nterm), or a mutant PDZD11 lacking the first 24 residues (P11-Δ24). Ponceau S-stained blots show baits (dashed black rectangles).

We generated antibodies against the C-terminal regions of PLEKHA5 and PLEKHA6 (Fig. S1A, orange boxes) and validated their specificity by immunoblotting (IB) and immunofluorescence (IF) analysis of cells either overexpressing (Fig. S2A,B) or lacking (Fig. S2C) the respective proteins. PDZD11 was detected in immunoprecipitates of PLEKHA5 and PLEKHA6, indicating the formation of a complex in cells (Fig. 1B). Moreover, in agreement with the results of the Y2H screen, GST-fusion proteins comprising the tandem N-terminal WW domains, but not the C-terminal domains of PLEKHA5 and PLEKHA6, interacted with full-length PDZD11 (Fig. 1C). Finally, full-length PLEKHA5 and PLEKHA6 interacted with GST baits comprising either full-length PDZD11 (Fig. 1D) or the first 30 residues of PDZD11, but not with PDZD11 lacking the first 24 residues (Fig. 1E), demonstrating that the WW domains of PLEKHA5 and PLEKHA6 interact with the N-terminal proline-rich region of PDZD11, similarly to PLEKHA7 (Rouaud *et al*., 2020).

### PLEKHA5, PLEKHA6 and PLEKHA7 show distinct localizations in cells and tissues and define cytoplasmic and microtubule-associated, lateral and junctional pools of PDZD11 in cultured cells

The cellular functions and localizations of PLEKHA5 and PLEKHA6 are not known. To determine whether the pattern of distribution of PLEKHA5 and PLEKHA6 is consistent with a role in ATP7A trafficking, we first examined their expression and localization in cells and tissues.

IB analysis showed distinct patterns of expression for each WW-PLEKHA in epithelial and non-epithelial cell lines (Fig. S3A and (Vasileva *et al*, 2017)). For IF analysis of endogenous proteins, we used epithelial cells from the collecting duct (mCCD) and the proximal tubule (MDCK) of the kidney, grown to full polarization either on Transwell filters or as cysts in Matrigel, and myeloblastic-derived Hap1 cells (Shah *et al*., 2018). mCCD cells express PLEKHA7 and PLEKHA6, while PLEKHA5 is not detectable by IB (Fig. S3A). In polarized mCCD cells, PLEKHA6 labeling was detected both at apical junctions, colocalizing with PLEKHA7, and along E-cadherin-labelled lateral contacts (Fig. 2A). Exogenously expressed GFP-tagged PLEKHA6 and PLEKHA7 are also detected at cell-cell contacts in mCCD cells and in bEnd.3 endothelial cells, while exogenous PLEKHA5 shows in addition a cytoplasmic fibrillar staining (Fig. S3B,C). Co-expression of PDZD11-HA with GFP-tagged PLEKHA6 and PLEKHA7 in polarized mCCD cells enhanced the accumulation of PDZD11 and its WW-PLEKHA interactor either along lateral contacts (PLEKHA6, Fig. 2B) or apical junctions (PLEKHA7, Fig. 2C). Co-expression of PDZD11 with PLEKHA5 resulted in enhanced localization of both proteins in the cytoplasm and along lateral contacts (Fig. 2D), whereas expression of PDZD11 alone resulted in its detection at apical junctions, lateral contacts and in the cytoplasm of mCCD cells (Fig. 2E).

**Figure 2.**
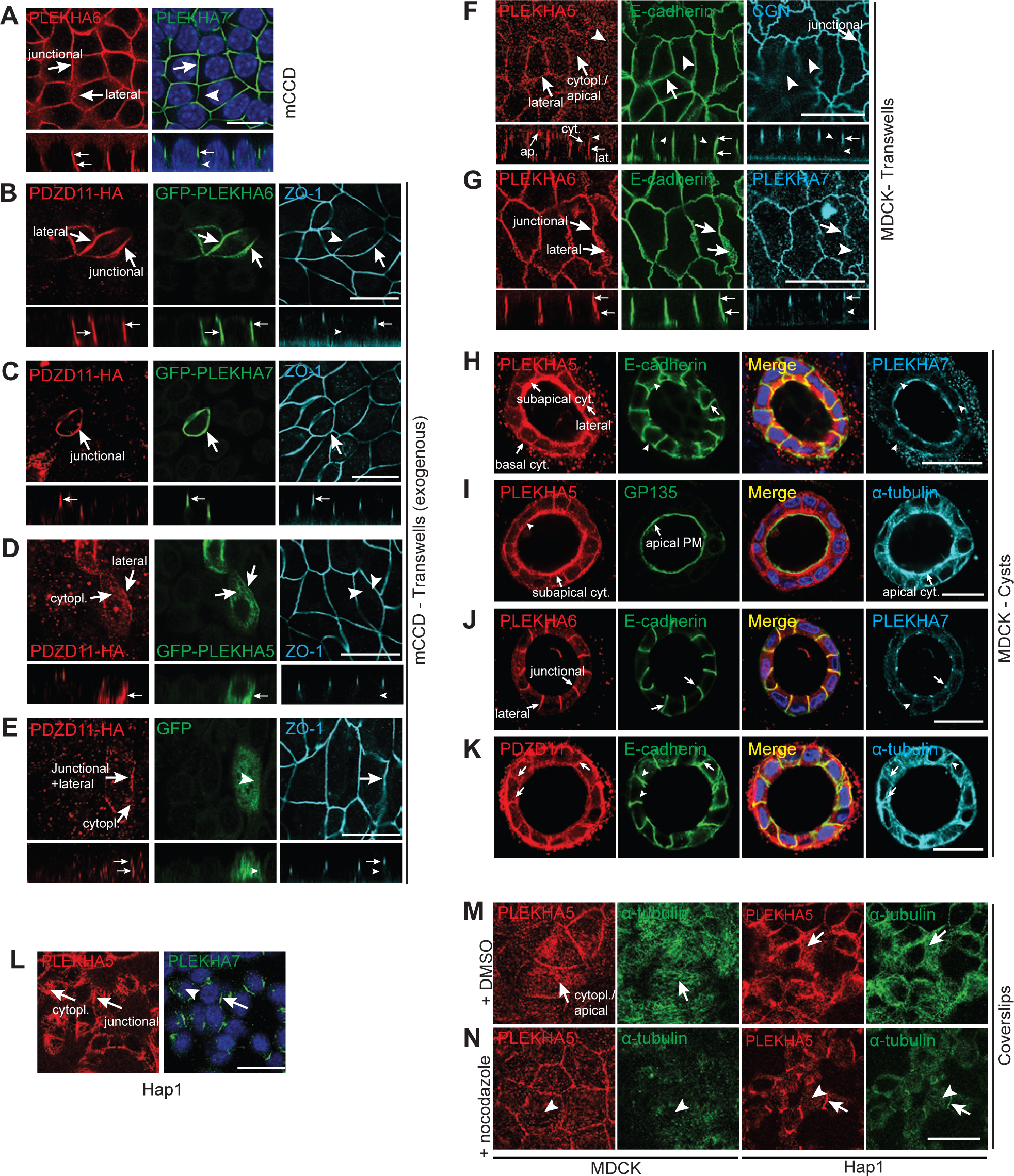
PLEKHA5 and PLEKHA6 are localized along the basal, apical and lateral plasma membranes in cultured cells and recruit PDZD11. (A, F-N) IF localization of endogenous PLEKHA5, PLEKHA6 and PLEKHA7 in cultured cells grown on different supports: (A) mCCD (Transwells); (F, G) MDCK (Transwells); (H-K) MDCK (cysts in Matrigel); (L) Hap1 (coverslips); (M,N) MDCK (left) and Hap1 (right) (coverslips), treated with either DMSO (M) or nocodazole (N). (B-E) IF localization of exogenous PDZD11-HA and GFP-tagged PLEKHA6 (B), PLEKHA7 (C), PLEKHA5 (D), and GFP alone (E, negative control) in mCCD cells (Transwells). For cells on Transwells, Z sections taken at the horizontal middle positions are shown below XY images. PLEKHA7, ZO-1 or cingulin (CGN) are used as markers for zonular apical junctions (ZA, TJ). Lateral contacts are labelled with E-cadherin. Arrows indicate labeling, arrowheads indicate low/undetectable labeling. Junctional, basal, cytoplasmic and lateral pools are indicated in epithelial cells. Bars= 20 µm.

MDCK cells express all three WW-PLEKHAs as detected by IB (Fig. S3A and (Pulimeno *et al*, 2010)). In MDCK cells grown on Transwells PLEKHA5 colocalized with E-cadherin at lateral contacts, in the submembrane cytoplasm near the apical and basal domains, and in the cytoplasm, but not at junctions (arrows and arrowheads, Fig. 2F), whereas PLEKHA6 was localized both at apical junctions, colocalizing with PLEKHA7, and along lateral contacts, colocalized with E-cadherin, but not in the cytoplasm (Fig. 2G, Fig. S3B). In MDCK cysts, PLEKHA5 was detected along the lateral membrane, colocalizing with E-cadherin, and in the cytoplasm near basal and apical plasma membranes (arrows, Fig. 2H). PLEKHA5 apical labeling was spatially distinct from that of the apical membrane marker GP135 but overlapped with tubulin labeling (arrows, Fig. 2I), indicating an association with apical submembrane structures associated with microtubules, rather than a juxtamembrane localization. PLEKHA6 was localized at junctions and along the lateral membrane (Fig. 2J), and PDZD11 was detected at junctions and lateral cell-cell contacts, and in the cytoplasm near basal and apical membranes, overlapping tubulin staining (Fig. 2K) similarly to PLEKHA5. Hap- 1 cells express PLEKHA5 and PLEKHA7, as detected by IB (Fig. S3A). In Hap1 cells PLEKHA5 was colocalized with PLEKHA7 at junctions (Shah *et al*., 2018) and also distributed also extensively along the plasma membrane and in a cytoplasmic fibrillar pattern (arrows and arrowheads, Fig. 2L). Since the localizations of PLEKHA5 and PDZD11 were similar to that of microtubules, we asked whether the integrity of the microtubule network controls PLEKHA5 localization. The cytoplasmic labeling of PLEKHA5 in both MDCK and Hap1 cells was dramatically reduced by treatment with the microtubule-depolymerizing drug nocodazole (Fig. 2M-N) indicating that cytoplasmic PLEKHA5 is associated with microtubules.

Next, we examined the expression and localization of WW-PLEKHAs in tissues. (IB) analysis showed expression of WW-PLEKHAs in different tissues and faster and slower migrating forms, suggesting the expression of differentially spliced isoforms and/or post-translational modifications (see also (Pulimeno *et al*., 2010)) (Fig. S3D). IF analysis of the kidney cortex showed strong PLEKHA5 labeling of glomeruli and weak diffuse labeling of apical and basal regions of tubular cells (arrows and arrowheads, Fig. S3E, top panels), whereas PLEKHA6 labeling was localized along the basal surface of tubular epithelial cells, and PLEKHA7 localized at epithelial apical junctions (arrows and arrowheads, Fig. S3E, bottom panels) (Pulimeno *et al*., 2010). In intestinal epithelial cells of the duodenum, PLEKHA5 was detected mostly in a subapical diffuse localization (Fig. S3F, top panel) whereas PLEKHA6 was detected both at junctions, colocalizing with PLEKHA7, and along basolateral surfaces (arrows, Fig. S3F, bottom panel). The localizations of PLEKHA5 and PLEKHA6 in intestinal epithelial cells were distinct from the localization of ATP7A, which is localized in the TGN (Fig. S3G) (Monty *et al*., 2005; Nyasae *et al*., 2007). We also tested whether WW-PLEKHAs are expressed in ATP7A-expressing cells in the brain. Strong expression of PLEKHA5 and PLEKHA7 were detected in blood vessels, marked by the endothelial marker PECAM- 1 (Fig S3H). ATP7A is highly expressed in pia matter adjacent to glial end-feet labeled with GFAP, and in the outer layer of the blood vessel (Fig. S3H). On the contrary, PLEKHA6 is not detected in brain blood vessel (Fig S3H, bottom panels). Expression of PLEKHA7 was broader than PLEKHA5, and in some cases, ATP7A colocalized with PECAM-1 and PLEKHA7 (Fig. S3H). The locus coeruleus (LC) contains the highest amount of copper among brain neurons and ATP7A plays an important role in metalate dopamine-β-hydroxylase activity to promote norepinephrine biosynthesis (Schmidt *et al*, 2018; Xiao *et al*, 2018). We tested whether PDZD11 and WW-PLEKHAs are co-expressed in these neurons. WW-PLEKHAs and PDZD11 were detected in LC neurons, however, ATP7A-labelled puncta did not overlap with WW-PLEKHAs and showed partial colocalization with PDZD11 in LC (Fig. S3I). Overall, PLEKHA7 distribution was broader than both PLEKHA5 and PLEKHA6, since labeling was detected in some neurons, brain blood vessels and choroid plexus, and shows a better overlap with ATP7A expression. On the other hand, although PLEKHA5 was immunolocalized in primary hippocampal neurons (Bayes *et al*, 2011; Pandya *et al*, 2017), our results on brain sections suggest a more abundant distribution in blood vessels. Finally, in primary cultures of cortical neurons, PLEKHA5 and PLEKHA6 labeling was detected in the cytoplasm of the neuronal cell body and was colocalized with β-tubulin-III along projections, whereas weak PLEKHA7 labeling was detected in the cytoplasm and nucleus (Fig. S3J, arrows). In summary, WW-PLEKHAs show different patterns of subcellular localization and tissue expression in cells and tissues that express ATP7A and do not colocalize with ATP7A under basal copper conditions.

### PDZD11 and WW-PLEKHAs are required for ATP7A localization at the cell periphery in response to elevated copper

To study the role of WW-PLEKHAs in the localization and trafficking of ATP7A we used available and new knock-out (KO) cell lines for either one or two WW-PLEKHAs, or for PDZD11, in the background of mCCD (Fig. S4A-C), MDCK (Fig. S4D-H) and Hap1 (Fig. S4I-K) cells. The KO of one WW-PLEKHA did not affect either the localization or levels of expression of the remaining WW-PLEKHA(s) (Fig. S5A, B, D, F). Moreover, KO of either PLEKHA5 or PLEKHA6 did not affect the localization of different components of junctions and lateral contacts, including nectin-3, ADAM10 and Tspan33, which interact with the PLEKHA7-PDZD11 complex (Fig. S5A, C, E) (Guerrera *et al*., 2016; Shah *et al*., 2018), or Tspan 15, which is localized laterally in epithelial cells (Fig. S5C,E) (Shah *et al*., 2018).

WT and KO clonal lines were cultured either as cysts in Matrigel or on Transwell filters, which allows optimal apicobasal polarization. In WT mCCD cells at basal copper levels ATP7A labeling was concentrated in a perinuclear location facing the apical membrane (apical perinuclear region), previously identified as the Trans-Golgi Network (TGN) (Greenough *et al*., 2004; Monty *et al*., 2005; Nyasae *et al*., 2007) (Fig. 3A, 3C and Fig. S6E, basal Cu). The same localization was observed at basal copper levels in mCCD cells KO for either PDZD11, or PLEKHA6, or PLEKHA7, or for both PLEKHA6 and PLEKHA7 (Fig. 3A, 3C and Fig S6A, C, E-I, basal Cu). Similarly, in both cells grown on Transwells and in cysts under basal copper conditions a perinuclear apical localization of ATP7A was observed in WT MDCK cells and in cells KO for either PLEKHA5 or PLEKHA6 (Fig. 4A, 4C and Fig. S7A-E, basal Cu). This indicates that neither PDZD11 nor WW-PLEKHAs are required for the localization of ATP7A in the TGN at basal copper levels in epithelial cells.

**Figure 3.**
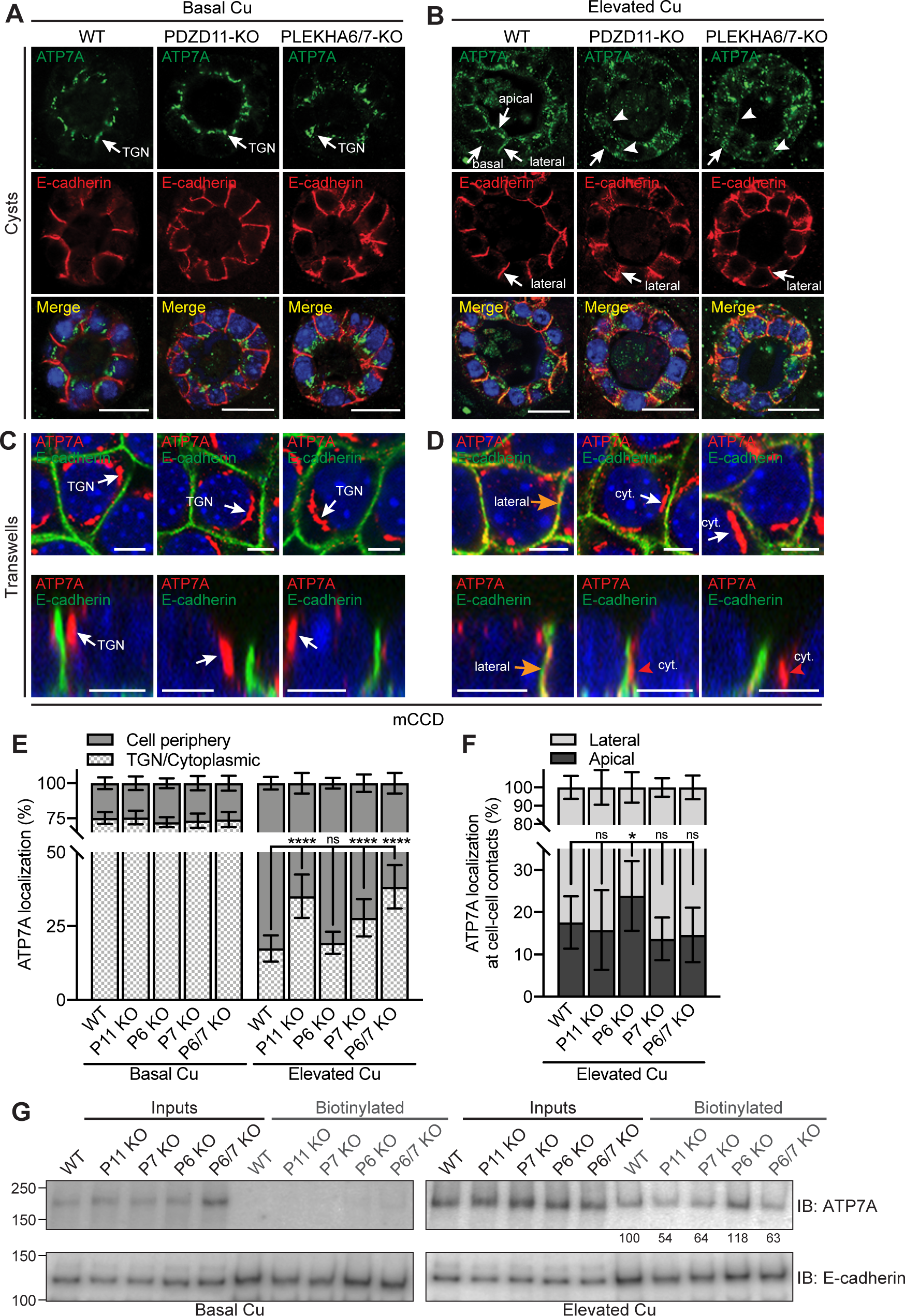
PDZD11, PLEKHA6 and PLEKHA7 are required for the correct targeting of ATP7A to the cell periphery of mCCD cells in elevated copper. (A-B) IF analysis of the localization of ATP7A in mCCD cysts. TGN= Trans-Golgi Network. Lateral, apical and basal ATP7A labeling in elevated copper are indicated by arrows in B. Arrowheads indicate low/undetectable labeling. Bars= 20 µm. (C-F) IF analysis (C-D) and quantifications (E-F) of ATP7A localization in mCCD cells gown on Transwells (see Figure S6 for images at lower magnification). (G) IB analysis of levels of biotinylated ATP7A in the indicated WT and KO lines under basal and high copper conditions. E-cadherin was used as positive control for basolateral labeling. Numbers below lanes indicate quantification of signal normalized to input, relative to WT cells. Orange arrows and red arrowheads in D indicate ATP7A labeling colocalized and non- colocalized with E-cadherin, respectively. Bars= 5 µm. Merge images indicate colocalization of ATP7A with E-cadherin (lateral contacts, green). Values are shown as mean ± SD. *n*=22-30 cells for Basal Cu and *n*=92-107 cells for Elevated Cu (E), *n=*18-23 cell-cell contacts (F). One-way ANOVA with post hoc Dunnett’s test (\**p<*0.05, \*\*\*\**p<*0.0001; ns, not significant).

**Figure 4.**
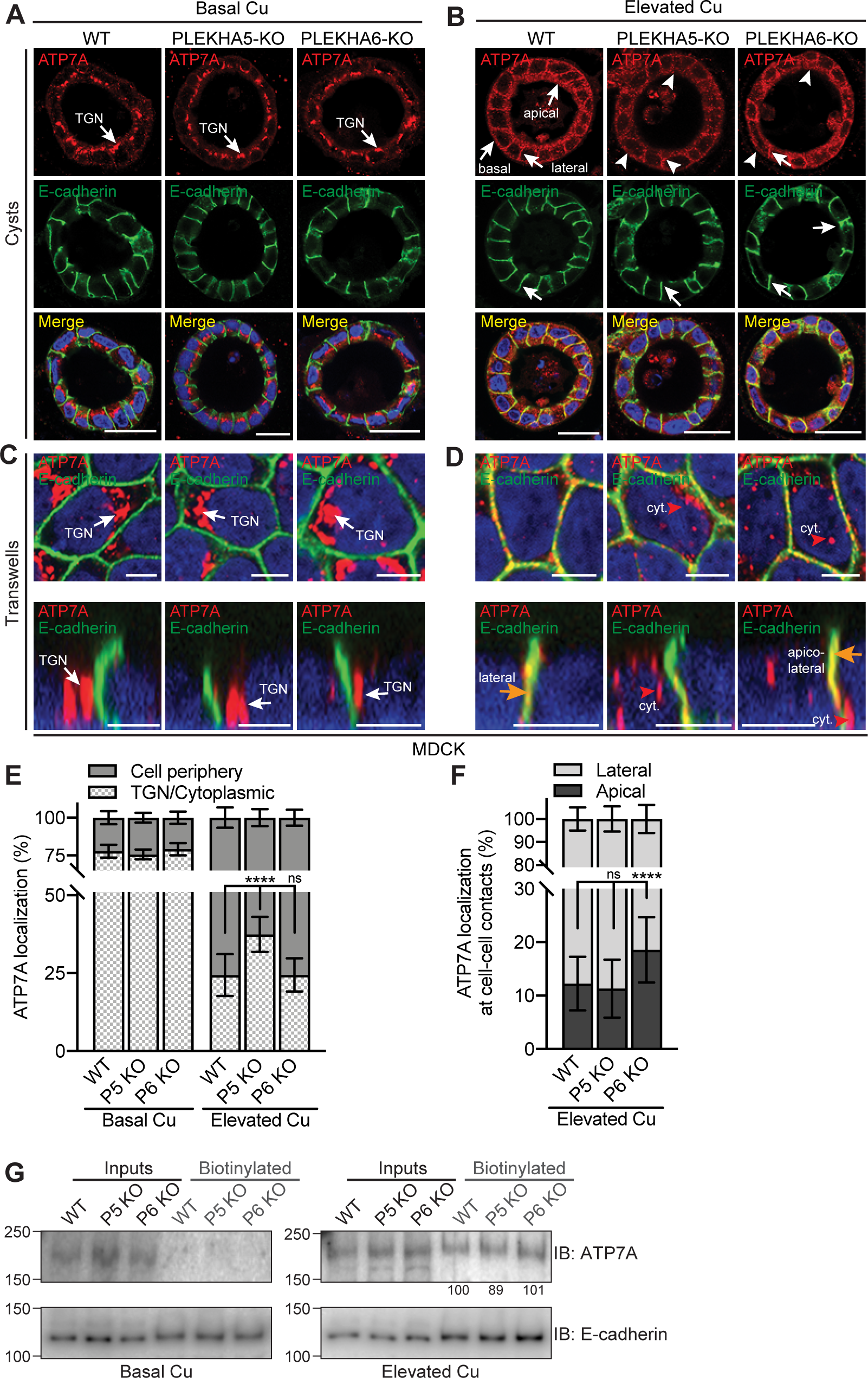
PLEKHA5 and PLEKHA6 are required for the correct targeting of ATP7A to the cell periphery of MDCK cells in elevated copper. (A-B) IF analysis of the localization of ATP7A in MDCK cysts. TGN= Trans-Golgi Network. Lateral, apical and basal ATP7A labeling in elevated copper are indicated by arrows in B. Arrowheads indicate low/undetectable labeling. Bars= 20 µm. (C-F) IF analysis (C-D) and quantifications (E-F) of ATP7A localization in MDCK cells gown on Transwells (see Figure S7 for images at lower magnification). (G) IB analysis of levels of biotinylated ATP7A in the indicated WT and KO lines under basal and high copper conditions. E- cadherin was used as positive control for basolateral labeling. Numbers below lanes indicate quantification of signal normalized to input, relative to WT cells. Orange arrows and red arrowhead in D indicate ATP7A labeling colocalized and non-colocalized with E-cadherin, respectively. Merge images indicate colocalization of ATP7A with E- cadherin (lateral contacts, green). Bars= 5 µm. Values are shown as mean ± SD. *n*=25- 31 (E, Basal Cu) and 90-105 cells (E, Elevated Cu), *n=*29-47 cell-cell contacts (F). One-way ANOVA with post hoc Dunnett’s test (\*\*\*\**p<*0.0001; ns, not significant).

Upon exposure to elevated copper, WT mCCD cells in cysts showed a massive redistribution of ATP7A labeling from the TGN to the cell periphery. Strong ATP7A labeling was detected linearly along lateral contacts, and near the apical and basal plasma membranes (arrows in Fig. 3B, WT). Instead, in mCCD cysts KO either for PDZD11 or for both PLEKHA6 and PLEKHA7, accumulation of ATP7A near the plasma membranes was disrupted (arrowheads in Fig. 3B, PDZD11-KO and PLEKHA6/7-KO).

In mCCD cysts KO for PLEKHA6 alone, the lateral accumulation of ATP7A was similar to WT cysts, whereas in cysts KO for PLEKHA7 alone the sub-membrane labeling was disrupted (Fig. S6B). In WT mCCD cells grown on Transwells, TGN labeling for ATP7A strongly decreased in elevated copper whereas labeling proximal to lateral E-cadherin was strongly increased (orange arrows in Fig. 3D, Fig. S6J). Instead, cells KO for either PDZD11, or both PLEKHA6 and PLEKHA7, or PLEKHA7 alone, showed decreased plasma membrane-associated ATP7A labeling colocalizing laterally with E-cadherin and increased cytoplasmic perinuclear staining (Fig. 3D, cyt. and red arrowheads, Fig. S6D, K, M, N, quantifications in Fig. 3E). In mCCD cells KO for PLEKHA6 the redistribution of ATP7A to the cell periphery was similar to that of WT cells (Fig. S6D, S6L, quantification in Fig. 3E), but labeling for ATP7A was shifted towards apical, rather than lateral cell-cell contacts (quantification in Fig. 3F).

In WT MDCK cysts exposed to elevated copper strong ATP7A labeling was detected along lateral contacts and near the apical and basal plasma membranes, similarly to mCCD cells (arrows in Fig. 4B, WT). The KO of PLEKHA5 resulted in a strong decrease in apical, basal and lateral ATP7A labeling in elevated copper, compared to WT cysts (arrowheads in Fig. 4B, PLEKHA5-KO). The KO of PLEKHA6 resulted instead in disorganized and reduced apical and basal labeling, but not as strong as that observed in PLEKHA5-KO cysts (arrow and arrowheads in Fig. 4B, PLEKHA6- KO). In MDCK cells grown on Transwells, KO of either PLEKHA5 or PLEKHA6 resulted in a significant increase in ATP7A labeling in the cytoplasmic space between the perinuclear TGN and the cell periphery (Fig. 4D and Fig. S7D-F, quantifications in Fig. 4E). Furthermore, in cells KO for PLEKHA6 lateral labeling for ATP7A was shifted apically (Fig. 4D, quantification in Fig. 4F).

Next, we asked whether either PDZD11 or WW-PLEKHAs are required for the retrograde trafficking of ATP7A to the TGN when the elevated copper levels are are depleted by treatment with the copper chelator bathocuproinedisulfonic acid (BCS). IF analysis revealed that both mCCD (Fig. S6O-S) and MDCK (Fig. S7G-I) WT and KO lines showed the same redistribution of ATP7A back to the TGN region after copper washout.

Finally, since previous work showed that copper induces increased basolateral plasma membrane levels of ATP7A, as measured by biotinylation (Greenough *et al*., 2004) we examined the role of WW-PLEKHAs and PDZD11 in controlling the basolateral surface levels of ATP7A. Basolateral surface proteins were biotinylated by addition of sulfo-NHS-SS-biotin to the basal chamber, isolated by affinity chromatography on streptavidin-coated beads, and lysates were analyzed by SDS-PAGE and IB using anti-ATP7A antibodies, and anti-E-cadherin antibodies as a positive control (Fig. 3G, Fig. 4G). No basolateral ATP7A was detected under basal copper conditions, whereas elevated copper resulted in detectable ATP7A (Fig. 3G, Fig. 4G), in agreement with previous results (Greenough *et al*., 2004). Confirming our IF analysis, KO of either PDZD11, or PLEKHA7, or both PLEKHA6 and PLEKHA7, but not PLEKHA6 alone, resulted in decreased levels of ATP7A at the basolateral surface of mCCD cells, when compared to WT (Fig. 3G). In MDCK cells, we observed a small decrease in the basolateral levels of ATP7A in PLEKHA5-KO, but not PLEKHA6-KO cells, when compared to WT (Fig. 4G).

In summary, these findings indicate that PDZD11 and WW-PLEKHAs are not required either for the TGN localization of ATP7A under basal copper conditions, for the copper-induced exit from TGN or for the retrograde traffic of ATP7A to the TGN after copper washout. However, they are required to promote and stabilize the localization of ATP7A at the cell periphery.

### PDZD11 and WW-PLEKHAs regulate intracellular copper homeostasis in response to elevated copper

We asked whether the altered localization of ATP7A in cells KO for either PDZD11 or WW-PLEKHAs correlates with changes in copper homeostasis. To this purpose, we used the copper Fluor-4 (CF4) probe (Fig. 5A, E), which, in combination with the control Copper Fluor-4-Sulfur-2 (Ctrl-CF4-S2) sensor that is insensitive to copper changes (Fig. 5B, F) provides a measure of intracellular labile copper (Xiao *et al*., 2018).

**Figure 5.**
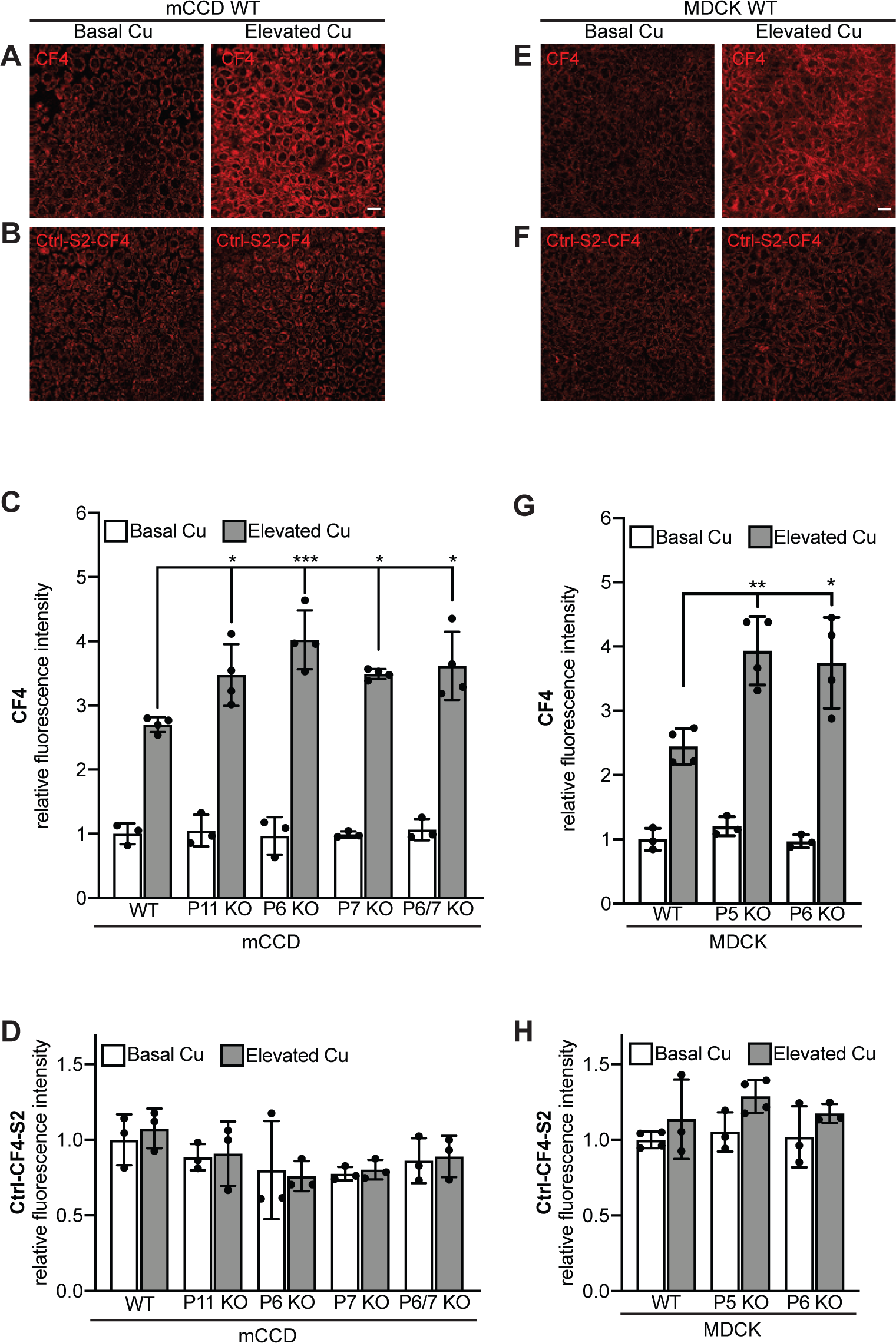
PDZD11 and WW-PLEKHAs are required for control of labile copper in mCCD and MDCK cells exposed to elevated copper. Fluorescence images (A-B, E-F) and quantification of fluorescence (C-D, G-H) of either mCCD (A-D) or MDCK (E-H) cells loaded either with CF4 (A,C,E,G) or Ctrl-CF4-S2 (B,D,F,H) under conditions of basal copper (Basal Cu) and after treatment with high concentration of copper (Elevated Cu). Abbreviations for genotypes (C,D,G,H): P11=PDZD11; P5=PLEKHA5, P6=PLEKHA6; P7=PLEKHA7. In C, D, G and H, dots show replicates (*n*), relative to the mean of WT cells in basal copper, and bars represent mean ± SD. One-way ANOVA with post hoc Dunnett’s test (\**p<*0.05, \*\**p<0.01*, \*\*\**p<*0.001).

In agreement with the normal TGN localization of ATP7A, in basal copper conditions KO cells (mCCD and MDCK) showed intracellular labile copper levels similar to those of WT cells, in both mCCD and MDCK lines (Fig. 5G). In contrast, exposure to elevated copper resulted in a 2.7-fold increase in intracellular labile copper in WT cells, but greater increases, between 3.4-fold and 4.1-fold in mCCD cells KO for either PDZD11, PLEKHA6, PLEKHA7 or both PLEKHA6 and PLEKHA7 (Fig. 5A-D). Similarly, in WT MDCK cells, elevated copper resulted in a 2.4-fold increase in intracellular labile copper, whereas in cells KO for either PLEKHA5 or PLEKHA6, the increase was 3.9- fold and 3.7-fold, respectively (Fig. 5G). The absence of significant differences in Ctrl-CF4-S2 fluorescent signals between WT and KO cells, either in basal or elevated copper environments, indicated a comparable loading of the probes in all conditions (Fig. 5D, H). The statistically significant increase in intracellular labile copper levels in KO cells exposed to elevated copper indicates that WW-PLEKHAs and PDZD11 regulate copper homeostasis.

### WW-PLEKHAs promote PDZD11 binding to ATP7A

Since PDZD11 binds both to the C-terminus of ATP7A and to the WW domains of WW- PLEKHAs, we investigated whether WW-PLEKHAs modulate ATP7A-PDZD11 interaction using a trimolecular GST pulldown assay. We used as preys GFP-tagged constructs encoding either the C-terminus of ATP7A (residues 1462-1500) or the same construct lacking the PDZ-binding motif (residues 1462-1496), or GFP alone (negative control) (Fig. 6A). As additional (third) proteins, we used HA-tagged constructs of PLEKHA5, PLEKHA6 and PLEKHA7 (Fig. 6B). Either GST-PDZD11 (Fig. 6C, 6F) or GST alone (negative control, Fig. 6D, 6G) were used as baits. IB analysis showed that the C-terminus of ATP7A but not GFP interacted with PDZD11, and this interaction increased significantly in the presence of WW-PLEKHAs (Figure 6C-E). In contrast, no interaction between PDZD11 and the C-terminus of ATP7A was observed when the PDZ-binding motif was deleted (Figure 6F-G). These results indicate that WW- PLEKHAs promote the interaction of the PDZ-binding motif of ATP7A with PDZD11.

**Figure 6.**
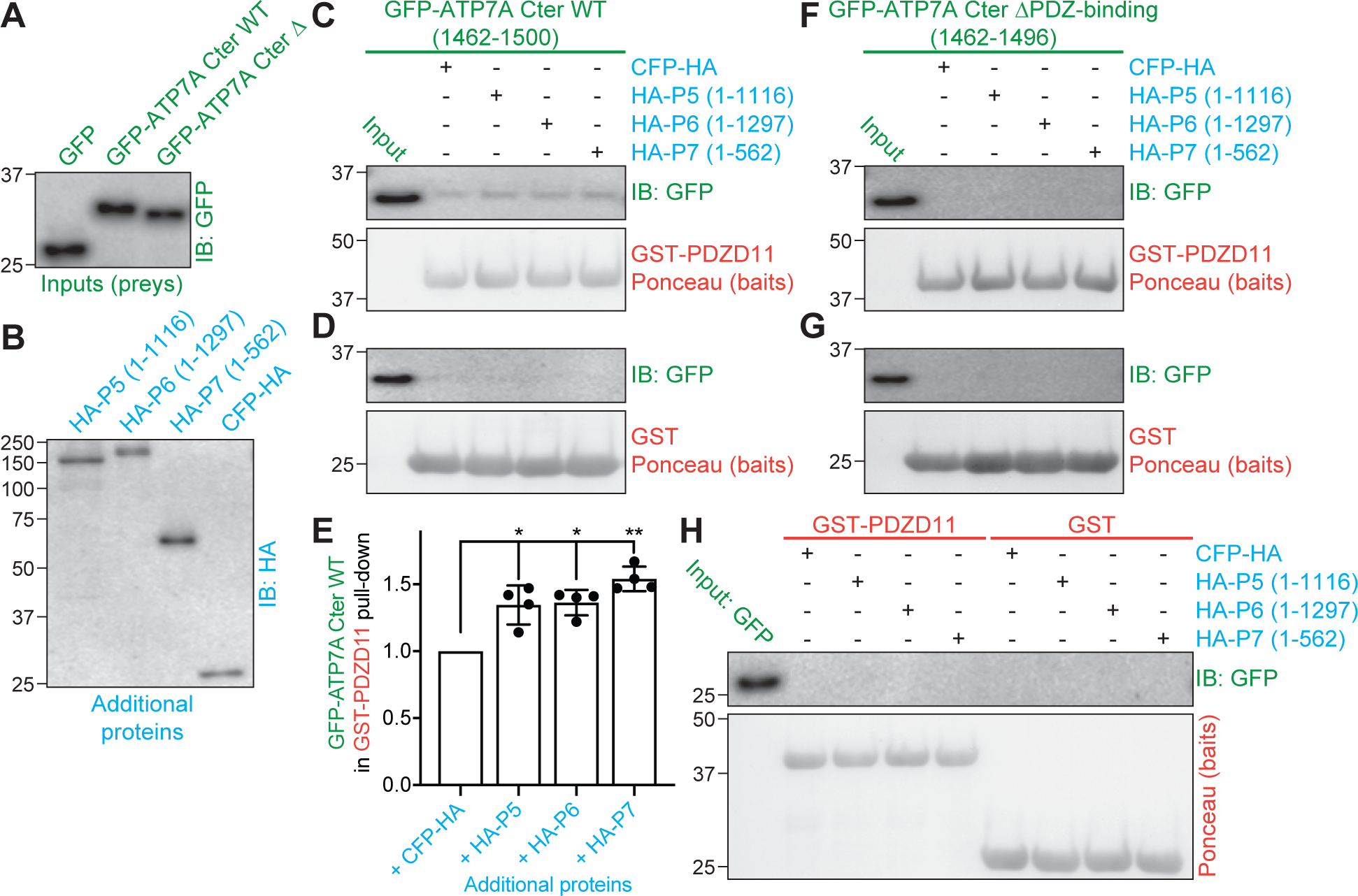
WW-PLEKHAs enhance the interaction of PDZD11 with the C-terminus of ATP7A. IB analysis using anti-GFP antibodies (C-D, F-H) and signal quantification (E) of GST pulldowns using either GST or GST-PDZD11 as baits (Ponceau S staining below IB show baits, in red). Preys (in green, normalization IB with anti-GFP shown A) were either GFP, GFP-tagged C-terminus of ATP7A WT (1465-1500) or lacking the C-terminal 4-residue PDZ-binding motif (GFP-ATP7A Cter ΔPDZ-binding). Pulldowns were carried out either in the presence of a HA-tagged third additional protein (in blue, normalization with anti-HA shown in B), either full length PLEKHA5 (P5), full-length PLEKHA6 (P6), N-terminal fragment of PLEKHA7 (P7), or CFP-HA (negative control). In E, dots show replicates (n) of GFP-ATP7A Cter WT pull-downs by GST-PDZD11 in the presence of WW-PLEKHAs relative to pull-downs in presence of the control additional protein (CFP-HA) (normalized to baits, see Material and Methods), and bars represent mean ± SD. Repeated measures (RM) one-way ANOVA followed by Dunnett’s multiple comparison test with +CFP-HA as reference (*p<0.05, **p<0.01).

## Discussion

The trafficking of ATP7A and the regulation of its localization is critical for the control of intracellular copper homeostasis, but little is known about the trafficking machinery that drives and stabilizes ATP7A at the cell periphery (Hartwig *et al*., 2019; Holloway *et al*., 2007; Holloway *et al*, 2013; La Fontaine & Mercer, 2007; Lutsenko *et al*., 2007; Polishchuk & Lutsenko, 2013; Skjorringe *et al*., 2017; Veldhuis *et al*, 2009). Here we identify PLEKHA5, PLEKHA6 and PLEKHA7, and their ligand adaptor PDZD11, as proteins involved in the copper-dependent localization of ATP7A at the cell periphery and in the maintenance of copper homeostasis.

PLEKHA5 and PLEKHA6 were first characterized as PH domain-containing proteins implicated in phosphoinositide signaling (Dowler *et al*., 2000), and genetic studies indicate that they participate in several developmental processes and diseases (Barbitoff *et al*, 2018; Cox *et al*, 2018; Daulagala *et al*, 2019; Fromer *et al*, 2014; Huang *et al*, 2020; Jamain *et al*, 2014; Jilaveanu *et al*, 2015; Liu *et al*, 2020; Shah *et al*, 2016; Spellmann *et al*, 2014; Tavano *et al*, 2018; Thapa *et al*, 2015; Wythe *et al*, 2011). However, little is known about the cellular localization and function of PLEKHA5 and PLEKHA6, and the molecular basis for the involvement of WW-PLEKHAs in physiological and pathological processes.

Here we show that the tissue distributions and subcellular localizations of PLEKHA5 and PLEKHA6 are distinct from PLEKHA7 and that WW-PLEKHAs and PDZD11 are expressed in essentially all cell types and tissues that express ATP7A. For example, in the adult mouse brain, ATP7A is expressed in endothelial cells of blood vessels (Qian *et al*, 1998), ependymal cells of choroid plexus in ventricles (Choi & Zheng, 2009; Kuo *et al*, 1997)), a subset of astrocytes (Kodama *et al*, 1991) and neurons (Iwase *et al*, 1996), and we detected WW-PLEKHAs in several of these cell types. PLEKHA5 appears less expressed than PLEKHA6 and PLEKHA7 in neuronal and epithelial tissues, and more expressed in vascular tissues. The subcellular localizations of WW- PLEKHAs are distinct, since PLEKHA5 is associated with microtubules and PDZD11 throughout the cytoplasm (see also (Zou & Cox, 2013)) and also detected near lateral and apical membranes, whereas PLEKHA6 is detected along basolateral membranes and apical AJ, and PLEKHA7 exclusively at apical AJ. The subcellular localization of ATP7A is highly sensitive to copper status, as low bioavailable copper levels direct ATP7A predominantly to the TGN, while high bioavailable copper levels target ATP7A to the cell periphery to facilitate copper efflux (Kaler, 2011). Importantly, ATP7A and WW-PLEKHAs are not colocalized under basal copper conditions, whereas they colocalize in elevated copper, suggesting a potential role in trafficking of ATP7A to the cell periphery upon elevated copper.

Our conclusion that the WW-PLEKHA-PDZD11-ATP7A interaction regulates ATP7A localization in elevated copper is based on the observation that the KO of either PDZD11 or WW-PLEKHAs, especially PLEKHA5 and PLEKHA7, correlated with decreased accumulation of ATP7A at the cell periphery, as detected both by IF analysis and biotinylation experiments. In PLEKHA6-KO cells ATP7A lateral labeling and biotinylated levels were not decreased, but lateral labeling shifted apically, suggesting that ATP7A is redundantly targeted to the lateral membrane by more than one WW-PLEKHA, but PLEKHA6 retains ATP7A laterally. Since PDZD11 is distributed in cellular pools associated with different WW-PLEKHAs, the redistribution of ATP7A towards the apical junction in PLEKHA6-KO cells may be driven by an increased proportion of PDZD11 bound to junctional PLEKHA7. The significance of the apical ATP7A labeling detected in MDCK cysts in elevated copper is unclear, since studies on mice show a redistribution of ATP7A from the TGN to basolateral but not apical regions of the plasma membrane in intestinal and kidney epithelial cells of mice exposed to elevated copper (Linz *et al*, 2008; Monty *et al*., 2005; Nyasae *et al*., 2007). Thus, future studies are required to explore the role of WW-PLEKHAs and PDZD11 in ATP7A traffic in vivo. In contrast, the KO of either PDZD11 or WW-PLEKHAs did not affect the localization of ATP7A in the TGN under basal copper and the copper-induced exit of ATP7A from the TGN. This is in agreement with the observation that ATP7A localization in the TGN in basal copper depends on sequences in the transmembrane domain 3 (Francis *et al*, 1998), and the copper-induced exit from the TGN depends on one CXXC metal binding site at the C-terminus (Goodyer *et al*, 1999; Lutsenko *et al*, 1997; Strausak *et al*, 1999), but not on the C-terminal sequences that interact with PDZD11 (Greenough *et al*., 2004; Stephenson *et al*., 2005). Finally, PDZD11 and WW-PLEKHAs did not control retrograde trafficking of ATP7A to the TGN after copper washout, in agreement with normal recycling of the ATP7A mutant lacking the PDZ-binding motif after restoration of basal copper levels (Greenough *et al*., 2004). Previous studies showed that internalization of ATP7A from the plasma membrane depends in part on clathrins and clathrin adaptors AP-1 and AP-2 and in part on clathrin-independent endocytic pathways (Holloway *et al*., 2013; Yi & Kaler, 2015). In summary, our results are consistent with a model whereby PDZD11 and WW-PLEKHAs help to target and stabilize ATP7A-containing vesicles to relevant domains of the plasma membrane (Fig. 7).

**Figure 7.**
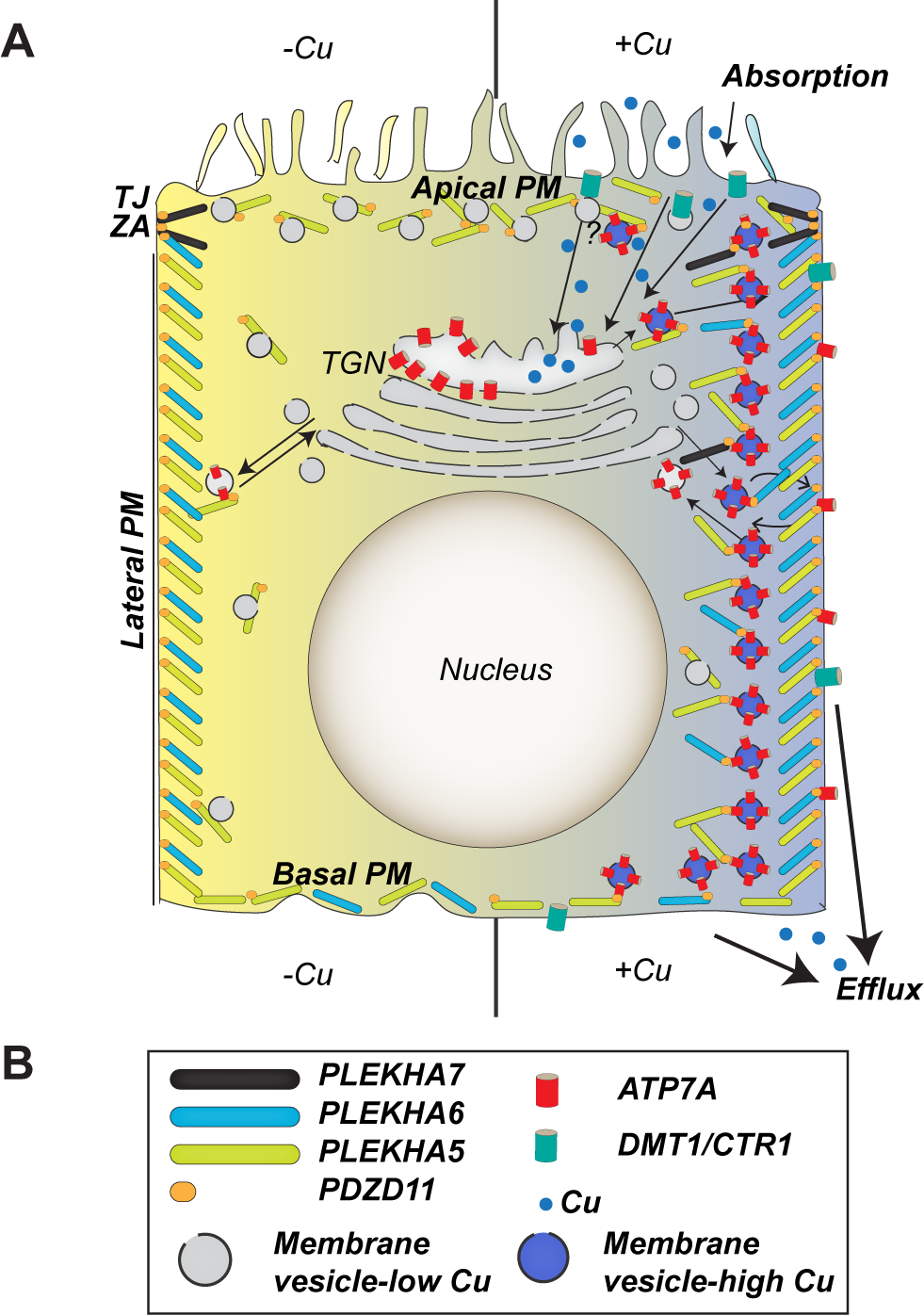
Model of functions of WW-PLEKHAs in ATP7A trafficking. Schematic model (A) and graphical legend (B) for the role of WW-PLEKHAs in the trafficking of ATP7A in absorptive polarized epithelial cells (e.g. MDCK), based on IF results shown in Fig. 3, Fig. 4, Fig. S6, Fig. S7. Apical, lateral and basal plasma membranes are shown. TJ and ZA denote tight junctions and *zonula adhaerens,* junctions that separate apical from lateral plasma membrane. PLEKHA7 is localized at the ZA. Under basal copper conditions (left), ATP7A-containing membrane vesicles cycle between the plasma membrane (PM) and Trans-Golgi Network (TGN), but ATP7A is mostly accumulated in the TGN. With elevated copper (right), ATP7A-containing membrane vesicles are trafficked to the cell periphery and are tethered to the apical lateral and basal PM by WW-PLEKHAs. In elevated copper, ATP7A loads copper (blue) into vesicles. Copper enters the cell through the copper transporters CTR1 and DMT1, and ATP7A is involved in copper efflux through circulation of ATP7A-containing membrane vesicles between TGN and basolateral plasma membrane. Other copper chaperones, transporters, adaptors and signaling proteins involved in copper homeostasis (La Fontaine & Mercer, 2007; Lutsenko *et al*., 2007; Polishchuk & Lutsenko, 2013) are not shown for the sake of simplicity.

We found that PDZD11 and WW-PLEKHAs regulate intracellular labile copper levels upon cell exposure to elevated copper. Although this could be simply explained by the reduced trafficking of ATP7A-containing vesicles to the plasma membrane, we did not observe a direct correlation between effect of KO on ATP7A localization and the increase in labile copper signal. This suggests that WW-PLEKHAs may regulate the activity of ATP7A independently of their effect on ATP7A localization, for example by modulating the recycling and dynamics of ATP7A-containing vesicles (La Fontaine *et al*., 1998; Pase *et al*, 2004; Petris *et al*., 1996), by anchoring vesicles to specific domains of the plasma membrane, or by maintaining a conformation of ATP7A that maximizes its function as a copper pump. It should be noted that the trafficking of ATP7A depends on actin and microtubule cytoskeletons (Cobbold *et al*, 2004; Cobbold *et al*., 2002), and PLEKHA7 was identified as a GTPase activating protein (GAP) for Rac1 and Cdc42 (Lee *et al*, 2017) and an indirect linker to microtubules (Meng *et al*, 2008), and we showed that PLEKHA5 associates with microtubules (see also (Zou & Cox, 2013)). Thus, another hypothesis is that WW-PLEKHAs regulate ATP7A trafficking by affecting the dynamics of the cytoskeleton. These questions, and the identification of additional components of the trafficking machinery for ATP7A, should be addressed by future studies.

We noted that WW-PLEKHAs promote the interaction of PDZD11 with the PDZ-binding motif of ATP7A. These results are in agreement with the idea that by interacting together WW-PLEKHAs and PDZD11 cooperatively promote the binding of the complex to their ligands (Guerrera *et al*., 2016; Rouaud *et al*., 2020; Shah *et al*., 2018). The WW domains of PLEKHA5 and PLEKHA6 interact with the N-terminal, Proline-rich sequences of PDZD11, similarly to PLEKHA7 (Guerrera *et al*., 2016; Rouaud *et al*., 2020), and the PDZ-binding motif of ATP7A is required for binding of the ATP7A C-terminus to the PDZD11-WW-PLEKHA complex. Thus, the mode of interaction of the WW-PLEKHA-PDZD11 complex with ATP7A is similar to what was described for the Ig-like adhesion molecules nectins (Guerrera *et al*., 2016) and distinct from that of Tspan33, which interacts directly with the first WW domain of PLEKHA7 (Rouaud *et al*., 2020). Collectively, our results indicate that PDZD11 links WW-PLEKHAs to the C-terminus of ATP7A and suggest that the WW-PLEKHA-PDZD11 complexes regulates the trafficking, membrane delivery and function of ATP7A and other transmembrane proteins. Future studies should address the mechanisms through which elevated copper acts as a transition metal signal to trigger the formation of the trimolecular ATP7A-PDZD11-WW-PLEKHA complexes (Ackerman & Chang, 2018; Chang, 2015).

## Experimental procedures

### Cell culture

Culture conditions for mouse cortical collecting duct cells (mCCD), Madin-Darby canine kidney cells (MDCKII Tet-off), mouse brain microvascular endothelial (endothelioma) cell line (bEnd.3), mouse heart endothelial cell line (H5V), human lung carcinoma cell line (A427), ciliated aortic mouse embryonic endothelial cells (meEC), human keratinocyte cell line (HaCaT) (Vasileva *et al*., 2017), human umbilical vascular endothelial cells (HUVEC) (Rouaud *et al*, 2019), haploid human cells (Hap1) (Popov *et al*, 2015), mouse mammary epithelial cells (Eph4), human intestinal carcinoma cells (Caco-2) (Spadaro *et al*, 2017), human embryonic kidney epithelial cells (HEK293T) (Rouaud *et al*., 2020) were described previously.

Cysts of mCCD and MDCK cells were obtained using the protocol of (Debnath *et al*, 2003). 40 µl of Matrigel (BD Biosciences, 354230) were added on glass coverslips in a 24-well plate and allowed to solidify for 30 min at 37°C. Cells were trypsinized, resuspended in SMEM medium (Sigma-Aldrich, M8167), pelleted by centrifugation (150x *g*, 3 min) and resuspended in 2 ml of SMEM to obtain a single-cell suspension. Cells were diluted to obtain 35000 cells/ml and mixed in a 1:1 ratio with Assay Medium 2x/Matrigel 4%/Epidermal growth factor (EGF) 10 ng/ml), and 400 µl were plated per well. MDCK cysts were grown up to 7 days, mCCD up to 14 days, replacing medium with fresh Assay Medium (1x) every 4 days.

Primary cultures of cortical neurons were obtained as described in (Chassefeyre *et al*, 2015). Cortices were dissected from E18.5 mouse embryos in HBSS (Invitrogen) containing Hepes 10 mM, streptomycin 10 µg/ml, penicillin 10 U/ml, then treated with 0.25% trypsin-EDTA for 10 min at 37°C and disrupted by 10 aspirations/ejections through a 1-ml micropipette tip followed by 10 cycles through a 200-µl tip. 400000 cells/well of dissociated cortical neurons were seeded in DMEM (Invitrogen) supplemented with 10% heat-inactivated horse serum in 6-well plates on 12-mm glass coverslips precoated overnight with 50 μg/ml poly-D-lysine (Thermo Fischer Scientific, A3890401) at 37°C. 20 hours after seeding, medium was changed to culture medium (Neurobasal (Invitrogen), B27 supplement 2%, sodium pyruvate 1 mM, L-Glutamine 2 mM, streptomycin 10 µg/ml, penicillin 10 U/ml), and 4 days after plating cytosine arabinoside (AraC, 5 µM) was added. Neurons were fed every 4 days with 500 µl of fresh culture medium containing AraC and fixed for immunofluorescence after 8 days of culture.

For immunofluorescence of copper-dependent ATP7A trafficking, MDCK or mCCD culture medium containing 315 µM CuCl2 (Sigma-Aldrich, C3279), diluted from aqueous 1500x stock solution, was added to the cells (Greenough *et al*., 2004), overnight before fixation in the case of cysts, 4 or 5 hours before fixation for MDCK or mCCD cells grown on Transwells, respectively. For copper washout, cells were treated with CuCl2 as described, washed once with culture medium, and then incubated for 4h at 37°C in culture medium containing 200 µM bathocuproinedisulfonic acid (Santa Cruz Biotechnology, Sc-217698) (copper chelating agent) and 50 µg/ml cycloheximide (protein synthesis inhibitor), with medium change every 80 min (Holloway *et al*., 2013).

To disrupt the microtubule network, cells were treated with 10 µM nocodazole (stock solution at 5 mg/ml in DMSO, Sigma-Aldrich, SML1665) for 2h at 37°C before immunofluorescence analysis. DMSO (maximal final concentration of 0.1%) treatment was used as negative control.

### Genome engineering

PLEKHA7 KO and PDZD11 KO mCCD and Hap1 cells were described previously (Guerrera *et al*., 2016; Popov *et al*., 2015; Shah *et al*., 2018). CRISPR/Cas9 gene editing technology was used to generate PLEKHA5 and PLEKHA6 KO cells, in wild type (WT) or PLEKHA7 KO background to obtain single or double KO cell lines, respectively. Genscript guide RNA (gRNA) designing tool (https://www.genscript.com/gRNA-design-tool.html) was used to determine the CRISPR target sequences (Table 1). After cloning the gRNAs into the BbsI site of Cas9 and GFP expressing px458 CRISPR plasmid (Addgene catalog no. 48138), specific background cells (Table 1) were transfected using Lipofectamine2000 (Invitrogen) and GFP-positive single-cell sorted as described previously (Guerrera *et al*., 2016; Shah *et al*., 2018). Single clones were further amplified and screened for KO using immunoblot and immunofluorescence analysis, before further validation by genotyping. Genomic DNA was purified using the DNeasy Blood and Tissue kit from QIAGEN (#69504) and the genomic locus surrounding the target region was amplified by PCR using specific primers (Table 1). Purified PCR products of Hap1 genotyping were subjected to Sanger sequencing (Microsynth, Switzerland), while products from mCCD and MDCK lines were first subcloned into the EcorI-HindIII site of pcDNA3.1(+)/myc-His to separate alleles before being sequenced. Due to missing regions in the genomic sequence of dog PLEKHA5, MDCK PLEKHA5 KO could not be genotyped.

**Table 1.**
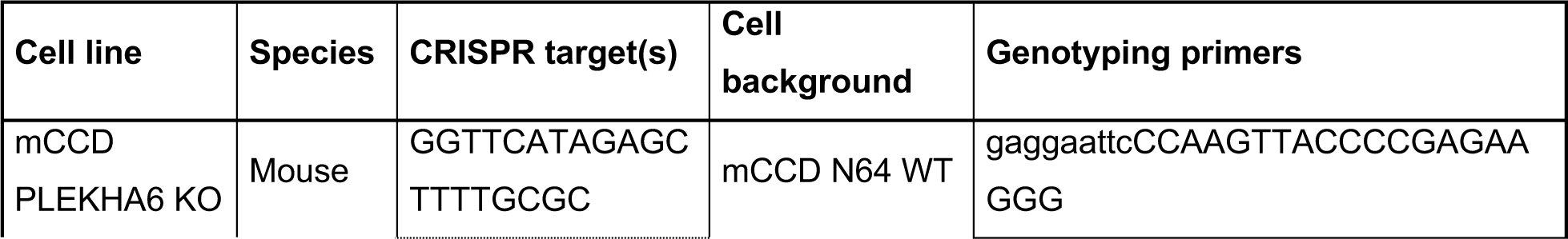
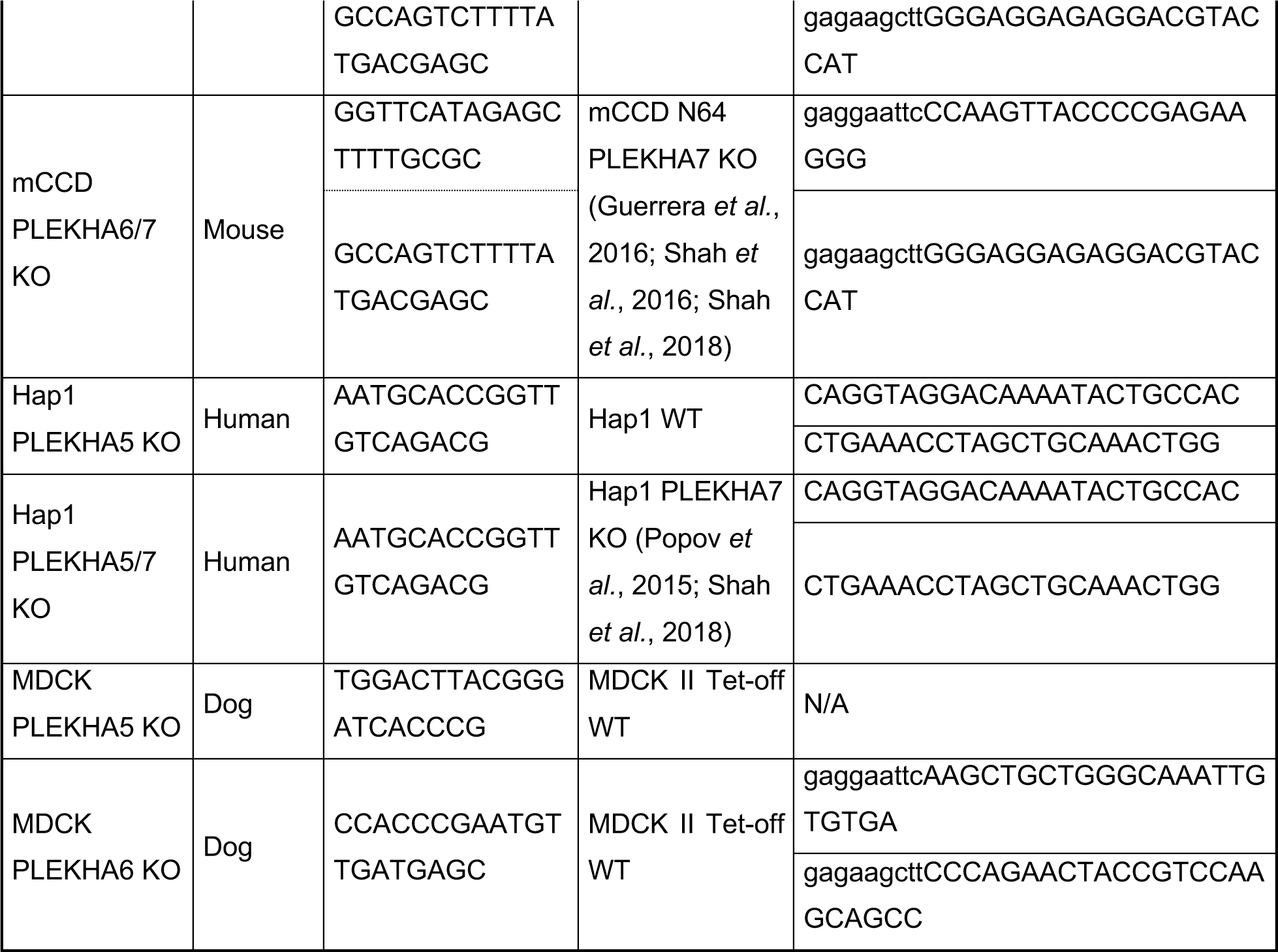
Genome engineering of PLEKHA5 and PLEKHA6 in mCCD, Hap1 and MDCK cells.

### Antibodies

The primary antibodies targeting the following proteins, raised in the detailed host species, were used at the indicated dilution for immunoblotting (IB), immunofluorescence (IF) or immunohistochemistry (IHC): rabbit PDZD11 (Rb29958, in-house (Guerrera *et al*., 2016), IB: 1/1000, IF,IHC: 1/100); rabbit PLEKHA7 (Rb30388, in-house (Pulimeno *et al*., 2010)}, IB: 1/5000, IF: 1/1000, IHC: 1/500); guinea pig PLEKHA7 (GP2737, in-house (Guerrera *et al*., 2016), IF: 1/500, IHC: 1/4000 or 1/300 for brain sections); mouse β-tubulin (32-2600, Thermo Fisher Scientific, IB: 1/3500); mouse α-tubulin (32-2500, Thermo Fisher Scientific, IF: 1/250); guinea pig α-tubulin (AA345 scFv-F2C, in house (Guerreiro & Meraldi, 2019), IF: 1/500); rabbit GFP (A-11122, Thermo Fisher Scientific, IB: 1/2000, IF: 1/200); mouse GFP (11814460001, Roche, IF: 1/100); mouse HA (32-6700, Thermo Fisher Scientific, IF: 1/150, IB: 1/1000); rabbit HA (sc-805, Santa Cruz, IF: 1/100); mouse myc (9E10, in-house, IB: 1/2); mouse E-cadherin (610181, BD Biosciences, IB: 1/5000, IF,IHC: 1/2500); mouse ZO-1 (33-9100, Thermo Fisher Scientific, IF: 1/1000); rat ZO-1 (R40.76, a kind gift from Prof. Daniel Goodenough (Harvard Medical School, USA), IF:

1/100); rabbit afadin (A0224, Sigma-Aldrich, IB: 1/8000, IF: 1/200); rabbit paracingulin (20893, in-house (Pulimeno *et al*, 2011), IB: 1/10 000, IF: 1/500); rabbit paracingulin (n.821, in-house (Guillemot *et al*, 2008), IF: 1/100); rabbit cingulin (C532, in-house (Cardellini *et al*, 1996), IF: 1/10 000, IB: 1/2000); mouse p120-catenin (8D11, a kind gift from Pr. A. Reynolds (Vanderbilt University, Nashville, USA) (Wu *et al*, 1998), IB: 1/2500, IF: 1/100); rabbit α-catenin (C2081, Sigma, IB: 1/8000); rabbit ß-catenin (C2206, Sigma, IB: 1/3500, IF: 1/500); rat nectin-3 (D084-3, MBL, IF: 1/100); rabbit ADAM10 (AB19026, Merck Millipore, IF: 1/300); mouse GP135 (3F2/D8, DSHB, IF:

1/5); mouse actin (MAB1501R, Merck Millipore, IB: 1/5000); mouse GFAP (G3893, Sigma-Aldrich, IF: 1/800); chicken GFAP (01-670-261, Invitrogen, IHC: 1/2000); mouse tubulin ß-III (801201, BioLegend, IF: 1/500); mouse ATP7A (sc-376467, Santa Cruz, IHC: 1/500); Armenian hamster PECAM-1 (MAB1398Z, Merck Millipore, IHC: 1/500); goat VE-cadherin (sc-6458, Santa Cruz, IF: 1/1000). The rabbit polyclonal antibody targeting ATP7A (RbCT78, IF,IHC: 1/500) (Steveson *et al*, 2003) was a kind gift from Betty Eipper (University of Connecticut Health Center, USA). The rat polyclonal antibodies targeting either PLEKHA5 (RtSZR129, IB: 1/1000, IF,IHC: 1/100) or PLEKHA6 (RtSZR127, IB: 1/1000, IF,IHC: 1/100) were generated by immunization of rats (Polyclonal Antibody Production, Eurogentec) with purified N-terminally GST-fused C-terminal fragments of human PLEKHA5 (NP_061885, aa 817-1116) or PLEKHA6 (XP_011507599, aa 971-1297) produced in BL21DE3 bacteria.

Secondary antibodies for IF, from Jackson ImmunoResearch and diluted at 1/300, were: anti-mouse (715-546-151), anti-rabbit (711-545-152), anti-rat (712-546-153) and anti-guinea pig (706-546-148) Alexa Fluor 488; anti-mouse (715-165-151), anti-rabbit (711-165-152) and anti-rat (712-166-153) Cy3; anti-guinea pig (706-605-148), anti-rabbit (711-605-152) and anti-goat (705-606-147) Alexa Fluor 647; anti-rat (712-175-153) and anti-mouse (715-605-151) Cy5. Additionally, for mouse brain IHC, secondary antibodies, diluted at 1/250, were: anti-Armenian hamster DyLightTM405 (127-475- 160, Jackson ImmunoResearch); anti-mouse Alexa Fluor Plus 555 (A32727, Thermo Fisher Scientific); anti-chicken Alexa Fluor Plus 488 (A32931, Thermo Fisher Scientific). Anti-mouse, anti-rabbit (1/20000, Promega, W4021 and W4011, respectively) and anti-rat (1/10000, Thermo Fisher Scientific, 62-9520) HRP-conjugated secondary antibodies were used for IB.

### Plasmids

Constructs that have been described previously include CFP-HA, full-length human PDZD11 (GFP-, GST- or -HA) and GST-fused WW (1-162) PLEKHA7 (Guerrera *et al*., 2016), GFP- and myc-tagged full-length (1-1121) and N-terminal region (1-562) of human PLEKHA7 (Paschoud *et al*, 2014), EGFP-tagged Tetraspanin-15 and Tetraspanin-33 (Shah *et al*., 2018), and GST-fused N-terminal (1-40) and Δ24 human PDZD11 (Rouaud *et al*., 2020). GFP-tagged WT (1462-1500) and ΔPDZ-binding (1462-1496) C-terminal regions of human ATP7A (NP_000043.4) were obtained by annealing of oligonucleotides and insertion in pcDNA3.1(-) plasmid previously modified to contain N-terminal GFP (pcDNA3.1-GFP) (Paschoud *et al*, 2011). Full-length sequences of human PLEKHA5 (NM_019012) and PLEKHA6 (XM_011509297) isoforms identified by Y2H screen were synthesized by Genscript (https://www.genscript.com/gene_synthesis.html) and amplified by PCR with appropriate oligonucleotides for subsequent cloning. GST-fusion of truncated sequences (PLEKHA5: WW (1-120), Cter (817-1116); PLEKHA6: WW (1-116), Cter (971-1297)) were cloned into pGEX4T1 (EcoRI/BamHI-NotI for PLEKHA5; EcoRI-NotI for PLEKHA6) for IPTG-inducible bacterial expression. For mammalian expression, full-length GFP-tagged PLEKHA5 (1-1116) and PLEKHA6 (1-1297) constructs were obtained by PCR and subcloned into pcDNA3.1(-) plasmid (NotI-KpnI for PLEKHA5, NotI-HindIII for PLEKHA6) previously modified to contain N-terminal GFP (pcDNA3.1- GFP) (Paschoud *et al*., 2011). N-terminally HA-tagged full-length PLEKHA5 and PLEKHA6 and N-terminal PLEKHA7 (1-562) were amplified by PCR using HA-containing forward primer before cloning into pcDNA3.1(-) (NotI-KpnI for PLEKHA5, NotI-HindIII for PLEKHA6 and PLEKHA7). Full-length PLEKHA5 and PLEKHA6, tagged with GFP in N-terminal and myc at the C-terminus, were made by PCR and subcloning into NotI-ClaI site of a pTre2Hyg plasmid already containing GFP-myc (Paschoud *et al*., 2014). All constructs were validated by sequencing (Microsynth, Switzerland).

### Yeast two-hybrid (Y2H) screen

The Y2H screen was carried out by Hybrigenics (France), with ULTImate Y2H screening technology. The full-length sequence of human PDZD11 (NP_001357103.1) was cloned in pB27 vector with N-terminal LexA, and this construct was used to screen a human placenta library (RP6), in the presence of 20 mM of 3-amino-1,2,4-triazole.

To perform bioinformatic analysis, the analyzed protein sequences corresponding to the human PLEKHA7 (NM_175058.4), PLEKHA5 (NM_019012.5) and PLEKHA6 (XM_011509297.2) were aligned using T-Coffee (version 8.93) from EMBL-EBI. WW and PH domains are SMART domains proposed by databases; coiled-coil domains were detected using NCoils (version 1) from Expasy; Proline-rich domains were obtained by a scan in the protein profile database PROSITE from Expasy. The two WW domains and PH domain of the three proteins were submitted to WebLogo 3.7.4 (Crooks *et al*, 2004) for graphical representation of the amino acid multiple sequence alignment of these regions.

### Cell transfection and immunofluorescence

For immunofluorescence (IF) staining, cells were seeded either on 6.5 mm/0.4 µm pore polyester 24-well tissue culture inserts (Transwell filters; Corning Costar #3470), or on 12-mm glass coverslips in 24-well plates. For Hap1 cells, coverslips were precoated with 0.01% Poly-L-lysine (Sigma-Aldrich, P4707) for 30 min at 37°C prior to plating. To study the localization of tagged proteins that are exogenously expressed, cells at 60 to 80% confluence were transfected one day after seeding, using either Lipofectamine2000 (Invitrogen) or jetOPTIMUS (Polyplus) following the manufacturer’s guidelines, and processed for IF 48-72h later. Cells on coverslips were washed twice with room temperature (RT) phosphate buffered saline (PBS) before methanol (pre-cold at -80°C) fixation during 8 min at -20°C. After three PBS washes, cells were permeabilized 5 min in PBS/Triton X-100 0.3% and blocked 20 min in blocking buffer (PBS/Gelatin 0.2%/Bovine serum albumin (BSA) 1%/Triton X-100 0.03% prior to incubation with primary antibodies, diluted in blocking buffer, either during 1h at RT or 16h at 4°C. Following three washes with PBS/Triton X-100 (0.3%) and 15 min of blocking, secondary antibodies and DAPI (1µg/ml), diluted in blocking buffer, were applied during 30 min at 37°C, before final washes with PBS/Triton X-100 (0.3%) (3 times) and PBS, and mounting with Fluoromount-G (Invitrogen). Alternative IF protocol, used when staining endogenous PLEKHA5 and PLEKHA6, consists in, after methanol fixation, washing the cells three times with PBS before a 30-min blocking step in PBS/donkey serum 1% and incubation overnight with primary antibodies diluted in serum incubation buffer (PBS/BSA 1%/donkey serum 1%/Triton X-100 0.3%) at 4°C. After three washes in PBS (15 min each), cells were incubated in secondary antibodies and DAPI (1 µg/ml), diluted in serum incubation buffer, during 30 min at 37°C, before final washes with PBS (three times, 15 min each) and mounting with Fluoromount-G (Invitrogen).

Cells grown on Transwells were fixed by 16h incubation in methanol at -20°C, followed by a 1-minute treatment with acetone pre-cooled at −20°C. Filters were excised manually using a razor blade and hydrated in IMF buffer (0.1% Triton X-100, 0.15 M NaCl, 5 mM EDTA, 20 mM HEPES, pH 7.5, 0.02% NaN3 as preservative) during 15 min at RT. After two washes with IMF, cells were blocked with IMF/donkey serum 1% for 30 min and then incubated with primary antibodies diluted in serum incubation IMF (IMF/BSA 1%/donkey serum 1%/Triton X-100 0.3%) overnight at 4°C. Three IMF washes (15 min each) were done prior to incubation with secondary antibodies (diluted in serum incubation IMF) for 2 hours at RT, and finals washes with IMF (four times, 15 min each). The filters were placed on glass slides, cells facing up, were mounted with Vectashield containing DAPI (VECTOR Laboratories) and covered by a glass coverslip.

Cysts were fixed with methanol and acetone mixed 1:1 for 11 min at -20°C before permeabilization with PBS containing 0.5% Triton X-100 (10 min at RT). Immunostaining was then performed as described previously (Spadaro *et al*., 2017).

Slides were imaged on a Zeiss LSM800 confocal microscope using a 63x/1.4NA oil immersion objective. Staining of nuclei with DAPI is colored in blue. Unless otherwise stated, scale bars correspond to 20 µm.

### Mouse tissues immunohistochemistry

Wildtype C57BL/6J mice were obtained from in-house breeding colonies. Mice were group housed on a 12:12 hour light-dark cycle at 22 °C with free access to food and water. All animal studies were approved by and performed according to the guidelines of the Animal Care and Use Committee of the University of Geneva (under authorization n. GE133/20) and of the University of California, Berkeley (under AUP- 2019-04-12038). To harvest epithelial tissues, mice were euthanized, tissues included in OCT medium and snap-frozen in liquid nitrogen-cooled isopentane. Frozen sections (5 µm) were air-dried, fixed with acetone at -20°C for 20 min and rehydrated in PBS. After 30 min of blocking in PBS/donkey serum 1%, sections were incubated with primary antibodies (overnight at 4°C) and secondary antibodies (1h at RT) diluted in PBS/BSA 1%/donkey serum 1%/Triton X-100 0.3%, each followed by three washings in PBS, and were finally mounted with Vectashield containing DAPI (VECTOR Laboratories) and covered by a glass coverslip. Sections were imaged on a Zeiss LSM800 confocal microscope using a 40x/1.3NA oil immersion objective. For brain IHC, mice were euthanized and immediately perfused with PBS and 4% formaldehyde in PBS. Brains were post-fixed in 4% paraformaldehyde for 24-48h and stored in 30% sucrose in PBS solution for 48 h for cryoprotection. Brains were embedded and mounted in Tissue-Tek OCT compound (Sakura finetek) and 20 µm sections were cut using a cryostat (Leica). Brain sections were rehydrated in PBS, permeabilized using PBST (0.3% Triton X-100 in PBS) for 30 min and incubated with blocking solution (5% normal goat serum or normal donkey serum in PBST) for 1h followed by primary antibody incubation overnight at 4°C. After washing in PBS, sections were incubated in corresponding fluorescently conjugated secondary antibodies (1/250) for 2 h at room temperature. After washing in PBS, sections were mounted with VECTASHIELD Antifade Mounting Medium (Vector Laboratories, H-1000). Fluorescence images were taken with a confocal microscope (LSM880 Confocal, Zeiss).

### Imaging quantifications

To quantify ATP7A localization between cytoplasmic/TGN and membrane-associated fractions (XY view), the integrated density of ATP7A signal in the cytoplasmic region (drawn inside the E-cadherin labeling with the polygon selection tool of FIJI) was divided by the integrated density of the signal in the entire cell area, i. e. comprising also the membrane-associated ATP7A staining, using E-cadherin to define this region with the polygon selection tool of FIJI. Quantification of ATP7A distribution at cell-cell contacts (XZ view) after copper treatment was done by calculating the zonular percentage of ATP7A signal, which was obtained by dividing the integrated density of the signal in the zonular region, using ZO-1 to delimit the area (polygon selection tool of FIJI), by the integrated density of the signal at the entire cell-cell contact area using E-cadherin to determine this region.

### Cell and tissue lysates, immunoblot analysis

Cell lysates were obtained in 500 µl of RIPA buffer (NaCl 150 mM/Tris-HCl 40 mM, pH 7.5/ EDTA 2 mM/glycerol 10%/Triton X-100 1%/sodium deoxycholate 0.5%/SDS 0.2%) supplemented with protease inhibitor cocktail (Thermo Fisher Scientific, A32965) from 10-cm dishes, followed by sonication (8 sec at 66% amplitude with a Branson sonifier). Solubilized proteins were clarified by centrifugation (15 min at 4°C, 13 000 rpm). Organ lysates were obtained by homogenization of the sample in 500 µl of lysis buffer A (Guillemot *et al*, 2012) using plastic micro-pestles. After 15 min of incubation on ice, samples were sonicated five times for 5 sec at 66% (Branson sonifier), before clarification by centrifugation (40 min at 4°C, 13 000 rpm).

Samples were mixed with SDS loading buffer and boiled 5 min at 95°C before SDS-PAGE separation at 4°C. Proteins were transferred onto nitrocellulose (0.45 µm) membrane (100V for 80 min or 70V for 180 min, at 4°C), and blots were blocked in Tris Buffered Saline (TBS)/Tween-20 0.1%/Low-fat milk 20% for 1 hr before incubation with primary antibody (diluted in TBS/Tween-20 0.1%/Low-fat milk 10%) followed by secondary HRP-labeled antibody (same buffer) for 1 hr, and finally chemiluminescence (ECL) revelation which was detected using Odyssey Imager (LI-COR). Numbers on the left of immunoblots correspond to sizes in kDa.

### Basolateral surface biotinylation

ATP7A levels along the basolateral surface was assessed by cell surface biotinylation (Greenough *et al*., 2004; Pase *et al*., 2004). 400 000 cells were grown on 24-mm transwells (Corning Costar #3450) for 7 days. For copper stimulation, culture medium containing 315 µM CuCl2 was added to the apical and basal chambers and incubated for 4-5h at 37°C. Biotin labelling and processing was performed at 4°C for 30 min, using Pierce Cell Surface Protein Isolation Kit (Thermo Fisher Scientific, 89881). Sulfo-NHS-SS-Biotin was dissolved in PBS supplemented with MgCl2 (0.5 mM) and CaCl2 (1 mM) (PBS^++^) and placed in the basal chamber, and PBS^++^ containing 315 µM CuCl2 was placed in the apical chamber. Biotin was quenched for 10 min and cells were scraped and transferred to a tube. After two washes with TBS (resuspension, centrifugation for 3 min at 500g and removal of supernatant), cells were lysed in lysis buffer (NaCl 150 mM/Tris-HCl 50 mM, pH 7.5/Triton X-100 1%/EDTA 5mM) containing protease and phosphatase inhibitor cocktail (Thermo Fisher Scientific, A32959) and sonicated two times for 4 sec at 45% (Branson sonifier). Lysates were incubated 30 minutes on ice, vortexing every 5 min for 5 sec, before centrifugation at 13 000 rpm for 15 min and transfer of the clarified supernatant to new tube. Biotinylated proteins were purified by overnight incubation with NeutrAvidin Agarose slurry from the kit. After incubation, beads were washed three times with lysis buffer, twice with high-salt buffer (NaCl 500 mM/ Tris-HCl 50 mM, pH 7.5) and once with no-salt buffer (Tris-HCl 10 mM, pH 7.5) before elution with sample buffer supplemented with DTT (200 mM) and urea (250 mg/ml) during 30 min at RT, vortexing every 5 min, and 15 min at 37°C. Samples were then analyzed by immunoblotting, along with inputs prepared in sample buffer containing DTT and urea (15 min of incubation at 37°C).

### Recombinant protein expression and Glutathione S-transferase (GST) pulldowns

For the production of GST-fused proteins, *E. coli* (BL21-DE3) were transformed by heat shock with pGEX4T1 constructs and expression was induced with 0.1 mM IPTG for 2h at 37°C. Bacterial pellets were snap frozen in liquid nitrogen before lysis in lysis buffer (PBS/Triton X-100 1%) supplemented with protease inhibitor cocktail (Thermo Fisher Scientific, A32965) and sonication five times at 55% (Branson sonifier). Cell debris were removed by centrifugation (13 000 rpm) for 15 min at 4°C, and GST-tagged baits contained in supernatants were normalized using Pierce Glutathione Magnetic Agarose Beads (Thermo Fisher Scientific, #78602) according to manufacturer’s protocol, followed by Coomassie staining of SDS-PAGE. Prey and additional (for tri- molecular pull-downs) proteins were expressed in HEK293T cells (2 300 000 cells in 10-cm dish) transfected with 10 µg of DNA using polyethylenimine (Polysciences, # 23966-2) and 48h later, after washing with PBS, lysing in CoIP buffer (NaCl 150 mM/Tris-HCl 20 mM, pH 7.5/Nonidet P-40 1%/EDTA 1 mM) supplemented with protease inhibitor cocktail (Thermo Fisher Scientific, A32965), applying sonication (8 sec at 66%, Branson sonifier) and centrifugation (15 min at 13 000 rpm, at 4°C). Prey and additional protein loadings were normalized by immunoblotting.

For GST pulldowns, 5 µg of GST-fused bait were coupled to Pierce Glutathione Magnetic Agarose Beads (Thermo Fisher Scientific, #78602), previously washed twice with equilibration buffer (Tris-HCl 125 mM, pH 7.4/NaCl 150 mM/DTT 1 mM/EDTA 1 mM), for 1h30 at RT. Following incubation and three washings with PBS/milk 2%/Nonidet P-40 1%, beads were incubated at 4°C overnight or for 2h with normalized preys HEK293T lysates. After three washings with CoIP buffer, proteins bound to the beads were eluated with 20 μl of SDS loading buffer boiled at 95°C for 5 min, before analysis by immunoblotting. Since lysates, and not purified proteins, were used as preys, it cannot be formally excluded that contaminating proteins from the HEK293T lysates may affect the results; however, it is unlikely that they are present in sufficiently high concentrations to affect results, as preys were significantly overexpressed.

Quantification of WT C-terminal ATP7A signal intensity in GST-PDZD11 pull-downs in the presence of WW-PLEKHAs was carried out in Image Studio Lite (LI-COR), normalized to bait signal (Ponceau S straining) and calculated relative to control additional protein (CFP-HA) value.

### Immunoprecipitation

Immunoprecipitation of proteins was carried out as described previously (Guerrera *et al*., 2016). Lysates were obtained from 10-cm dishes by rinsing cells with PBS and incubating them in 500 µl of CoIP buffer (NaCl 150 mM/Tris-HCl 20 mM, pH 7.5/Nonidet P-40 1%/EDTA 1 mM) supplemented with protease inhibitor cocktail (Thermo Fisher Scientific, A32965) for 10 min at 4°C. After sonication (8 sec at 66%, Branson sonifier) and centrifugation (15 min at 13 000 rpm, at 4°C), the supernatant was collected (cytoskeleton-soluble fraction). The pellet was resuspended in 50 µl of SDS buffer (SDS 1%/Tris-HCl 10 mM, pH 7.5/EDTA 2 mM/DTT 0.5 mM/PMSF

0.5 mM), sonicated 3 sec at a power of 15% (Branson sonifier), incubated 5 min at 95°C and clarified by centrifugation. Supernatant was brought to a volume of 500 µl with CoIP buffer and mixed with the cytoskeleton-soluble fraction to obtain the total cell lysate. 20 µl of Dynabeads protein G (or protein A for guinea pig anti-PLEKHA7) (Invitrogen) were coupled to antibodies (diluted in PBS/BSA 5%/Nonidet P-40 1%; 2 µl of pre-immune or immune serum for anti-PLEKHA7, 10 µl for anti-PLEKHA5 and -6) at 4°C for 90 min. After two washes with PBS/BSA 5%/Nonidet P-40 1%, beads were incubated overnight at 4°C with 100-120 µl of total cell lysate and then washed three times with CoIP buffer. Immunoprecipitates were eluted in 20 µl SDS loading buffer and boiled 5 min at 95°C, before analysis by SDS-PAGE and immunoblotting.

### Intracellular labile copper imaging

Cells were seeded on 35-mm glass-bottom fluorodishes (WPI, FD35-100) and incubated for 48hr until confluent. On the day of imaging, after 4 (MDCK) or 5 (mCCD) hours of incubation at 37°C (5% CO2) in fresh culture medium at basal or elevated (315 µM, CuCl2 mixed as aqueous solution) copper, cells were washed with Live Cell Imaging Solution (LCIS, Thermo Fisher Scientific, A14291DJ) before loading with either 1 µM CF4 (diluted in LCIS) or 1 µM Ctrl-CF4-S2 probe (Xiao *et al*., 2018) for 20 min at 37°C. Cells were rinsed twice with LCIS and imaged in LCIS at 37°C on a Zeiss LSM780 confocal microscope using a 40x/1.2NA water immersion objective, exciting the probe at 536 nm (HeNe543 laser) and collecting emission between 545 and 700 nm. 7-22 fields were acquired for each fluorodish. In FIJI, signal was adjusted to threshold to remove background areas and the mean intensity of the image (limited to threshold) was measured; the average of the 7-22 intensities was calculated for each fluorodish (replicate). GraphPad Prism 8 software was then used to normalize replicates to the mean of WT cells in basal copper conditions.

### Statistical analysis

Statistical significance was determined using GraphPad Prism 8 software; sample size (n), p-values (P) and statistical tests performed are specified in figure legends. Values are shown as mean ± standard deviation (SD). Significance has been determined as follows: *p<0.05, **p<0.01, ***p<0.001, ****p<0.0001. For multiple comparison tests, after ANOVA results showing significant difference, Dunnett’s test was used to compare every mean to control mean.

## Author Contributions

SS, IM, TX, AB, FF, and LJ conducted experiments. CJC provided reagents. SS performed statistical analysis. SS and SC designed experiments, analyzed the data, and wrote the paper.

## Acknowledgments

This work was supported by the Swiss National Science Foundation (31003A_172809) (to S.C), by the State of Geneva, and by grant NIH GM 79465 (to C.J.C.). We thank the Service Egalité of the Unversity of Geneva for a “Subside Tremplin” (to S.S.). We thank Arielle Flinois and Marine Laporte for protocols of cysts and neuronal culture, Betty Eipper and Scot Leary (University of Saskatchewan, Canada) for gift of anti- ATP7A antibodies, Quentin Devos and Thomas Di Mattia for informatic help.

## KEY RESOURCES TABLE

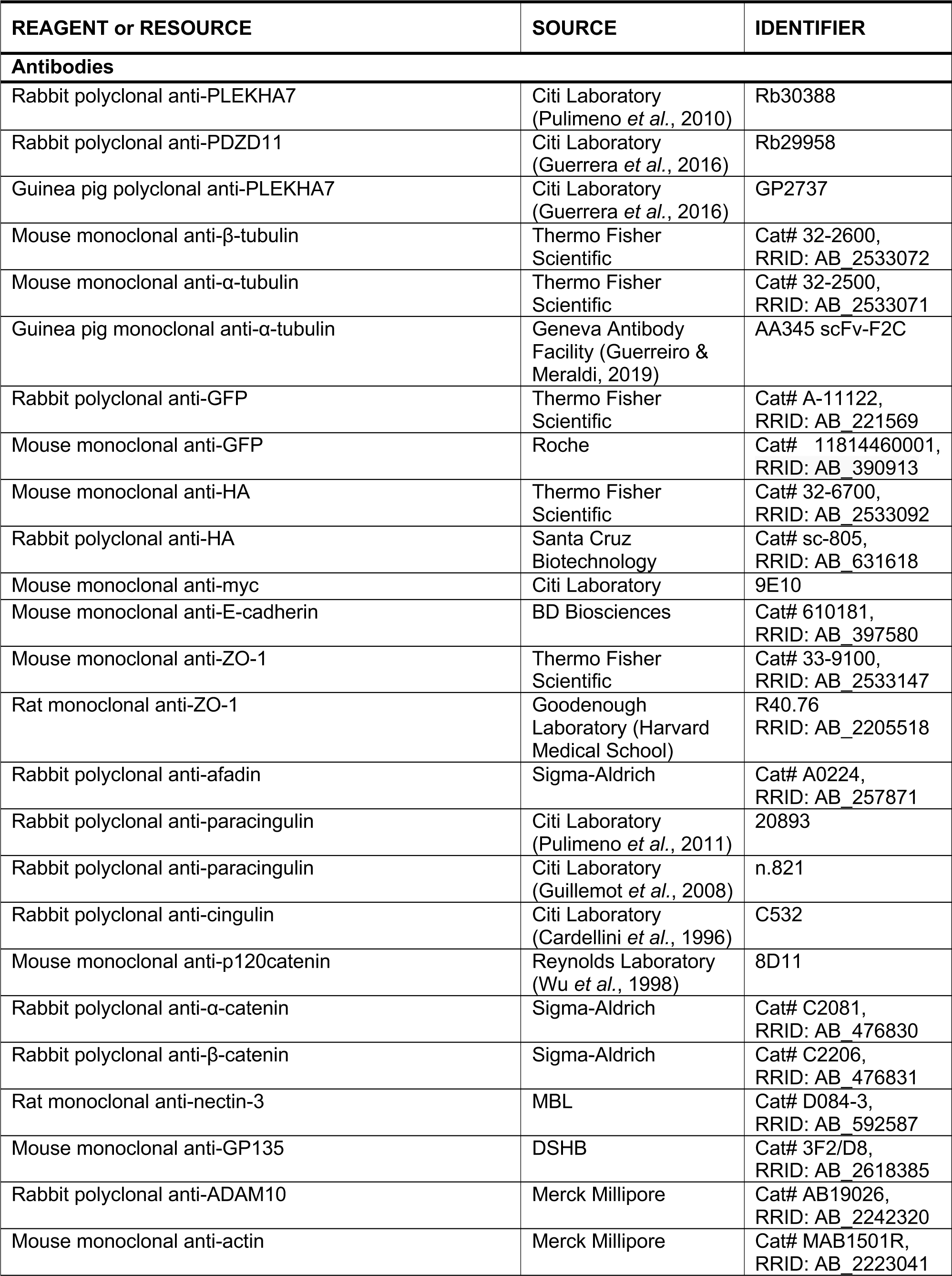
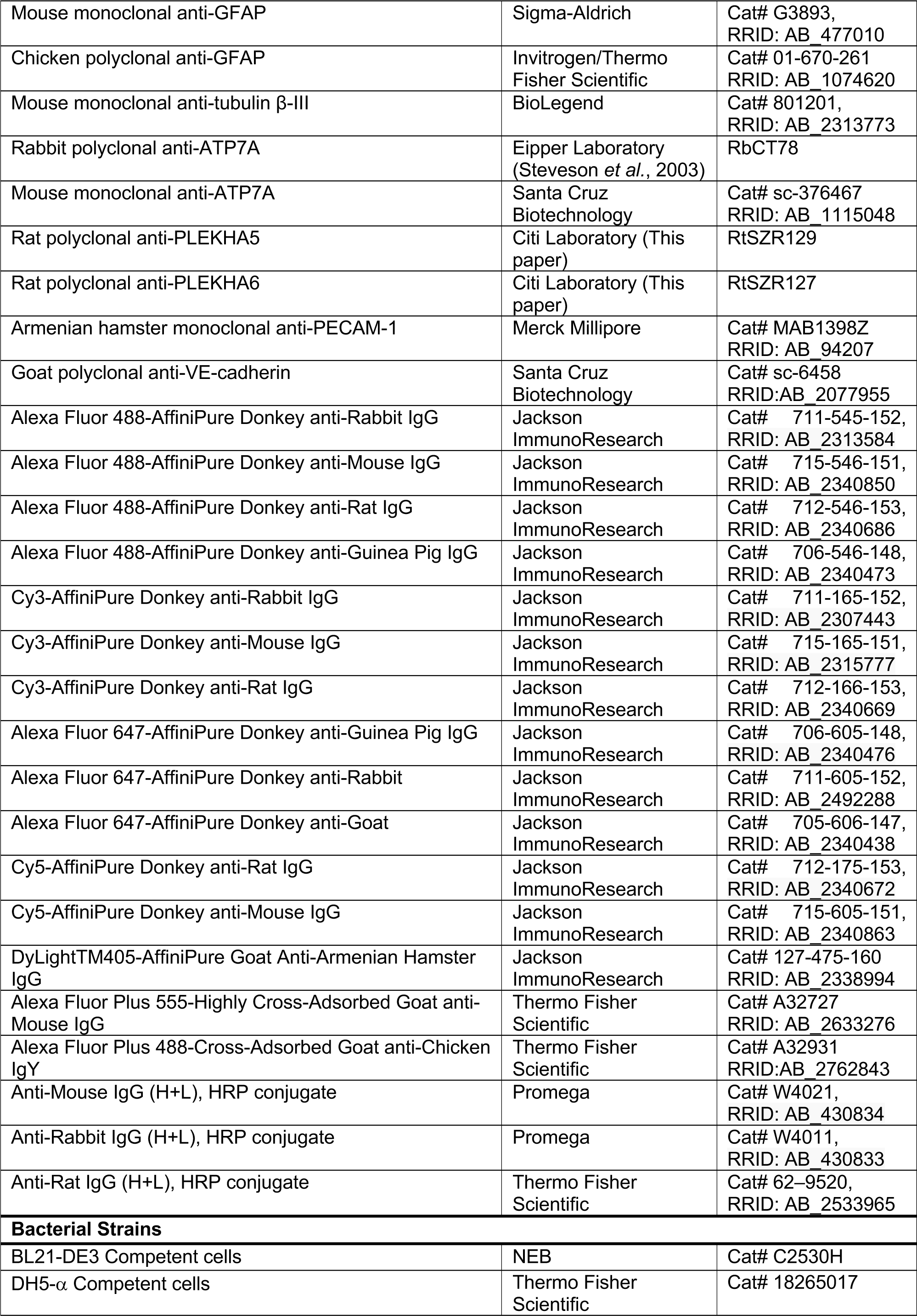
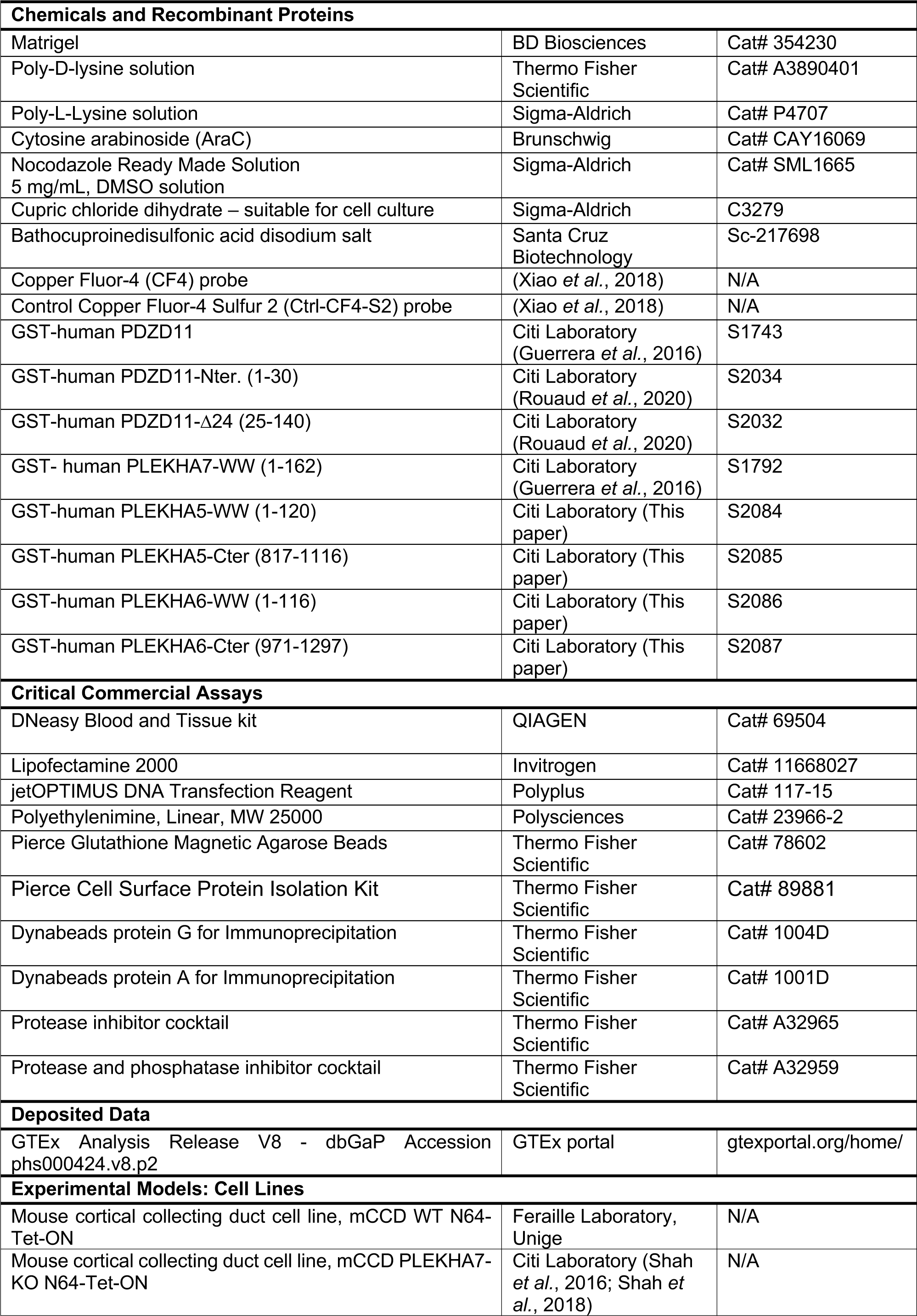
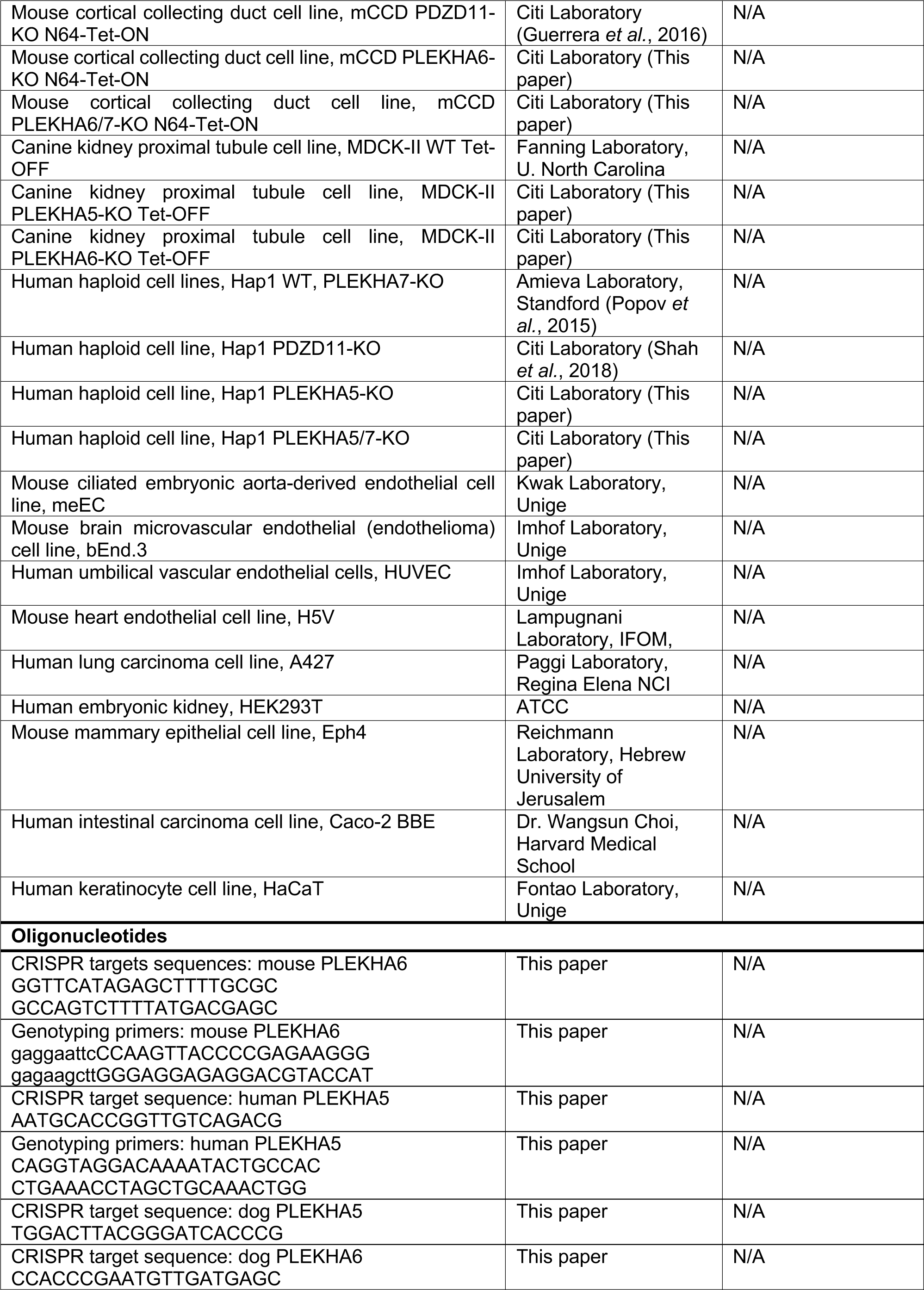
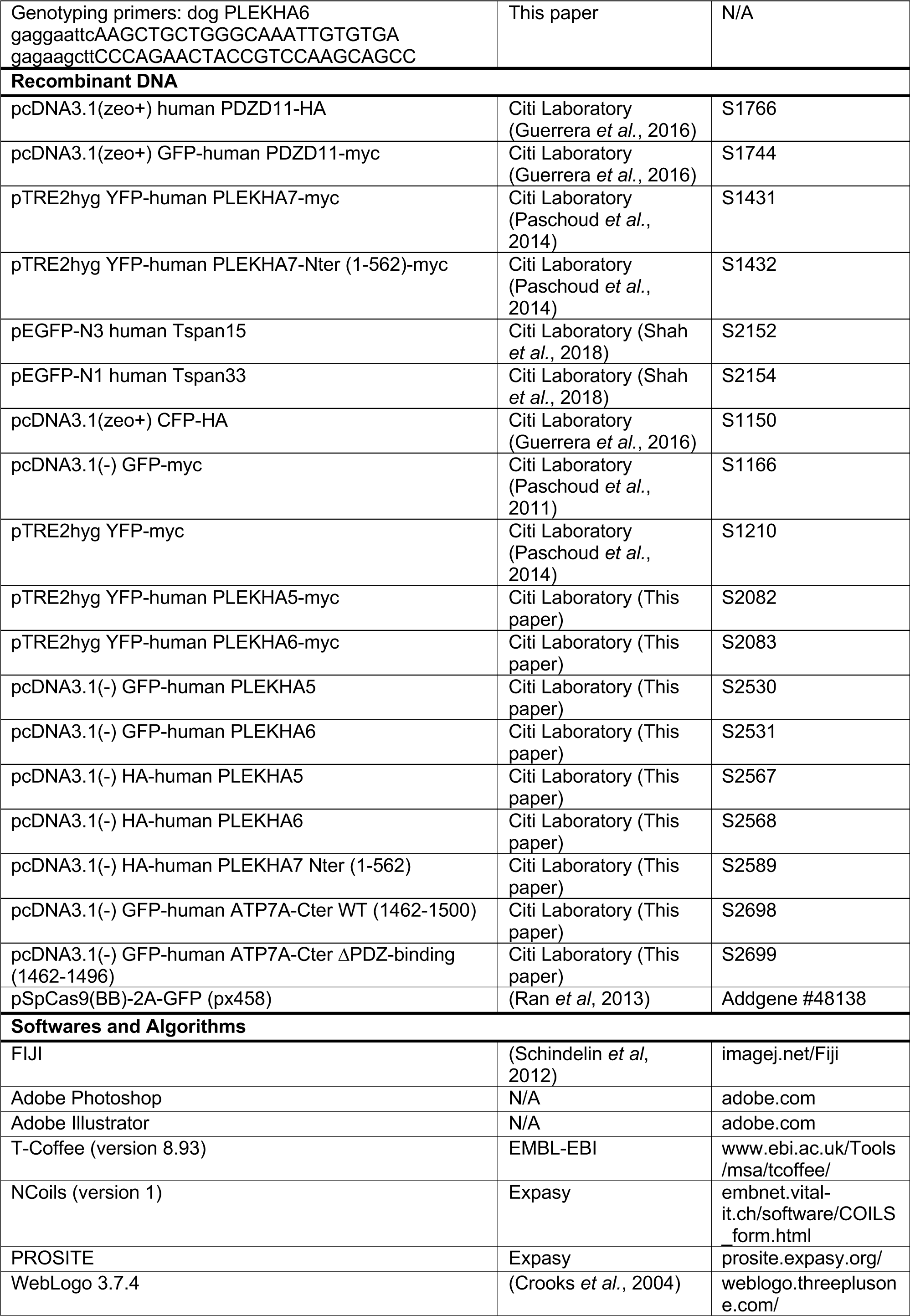
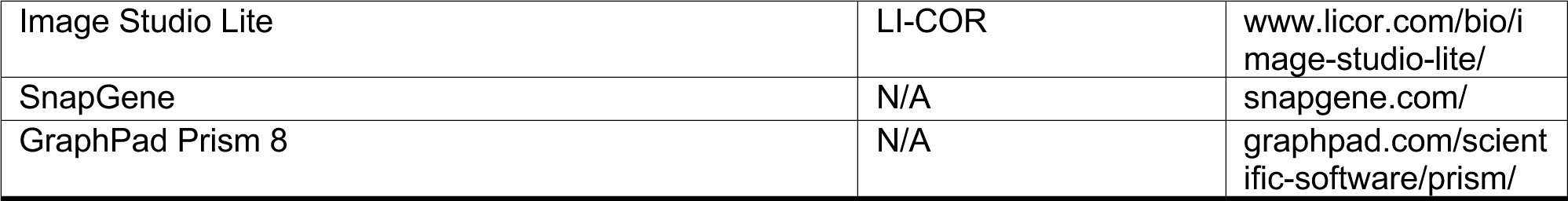

## SUPPLEMENTARY FIGURE LEGENDS

**Figure S1.**
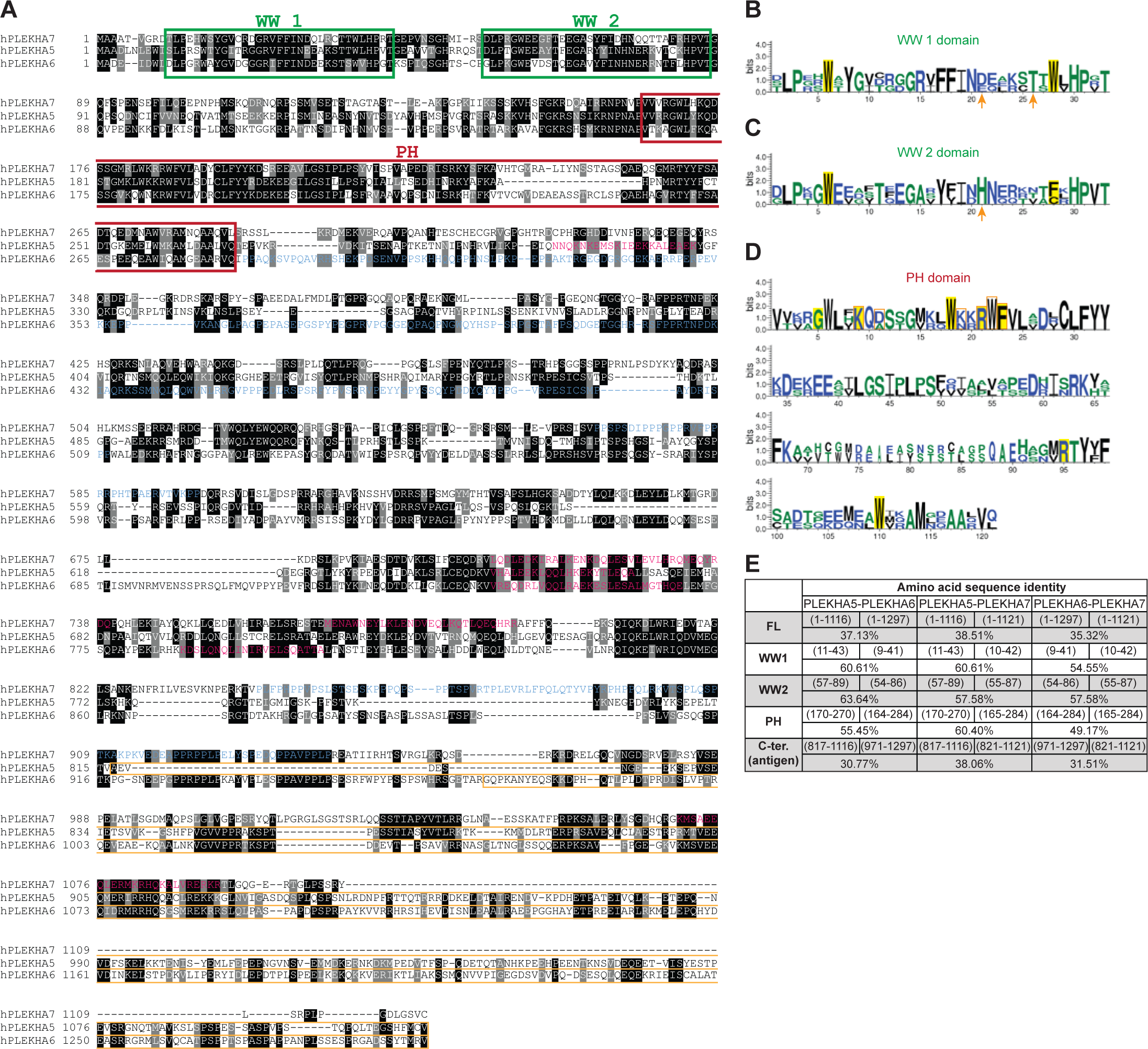
Sequence homology between PLEKHA5, PLEKHA6 and PLEKHA7. (A) Multiple sequence alignment of human PLEKHA5, PLEKHA6 and PLEKHA7 showing identical (black) and similar (grey) residues. WW and PH domains are in green and red boxes, respectively. Residues of coiled-coil and proline-rich domains are indicated in pink and blue, respectively. Orange boxes show regions used as antigens for generation of antibodies. (B-D) Weblogo diagrams of residue conservation in the first (B) and second (C) WW domains, and in the PH domain (D) of human PLEKHA5, PLEKHA6 and PLEKHA7. In B and C, signature residues of the WW domains are highlighted in yellow, and arrows point the amino acids forming the pocket for interaction with PDZD11 (Rouaud *et al*., 2020). In D, the residues that make up the putative PtdIns(3,4,5)P3-binding motif (PPBM) (Dowler *et al*., 2000; Isakoff *et al*, 1998)are highlighted in yellow, and the key amino acids for the PtdIns(3,4,5)P3 binding PH motif are squared in orange (Jungmichel *et al*, 2014)}. (E) Percentage values of amino acid sequence identity between full-length and domains (with indicated amino acid positions) of WW-PLEKHAs.

**Figure S2.**
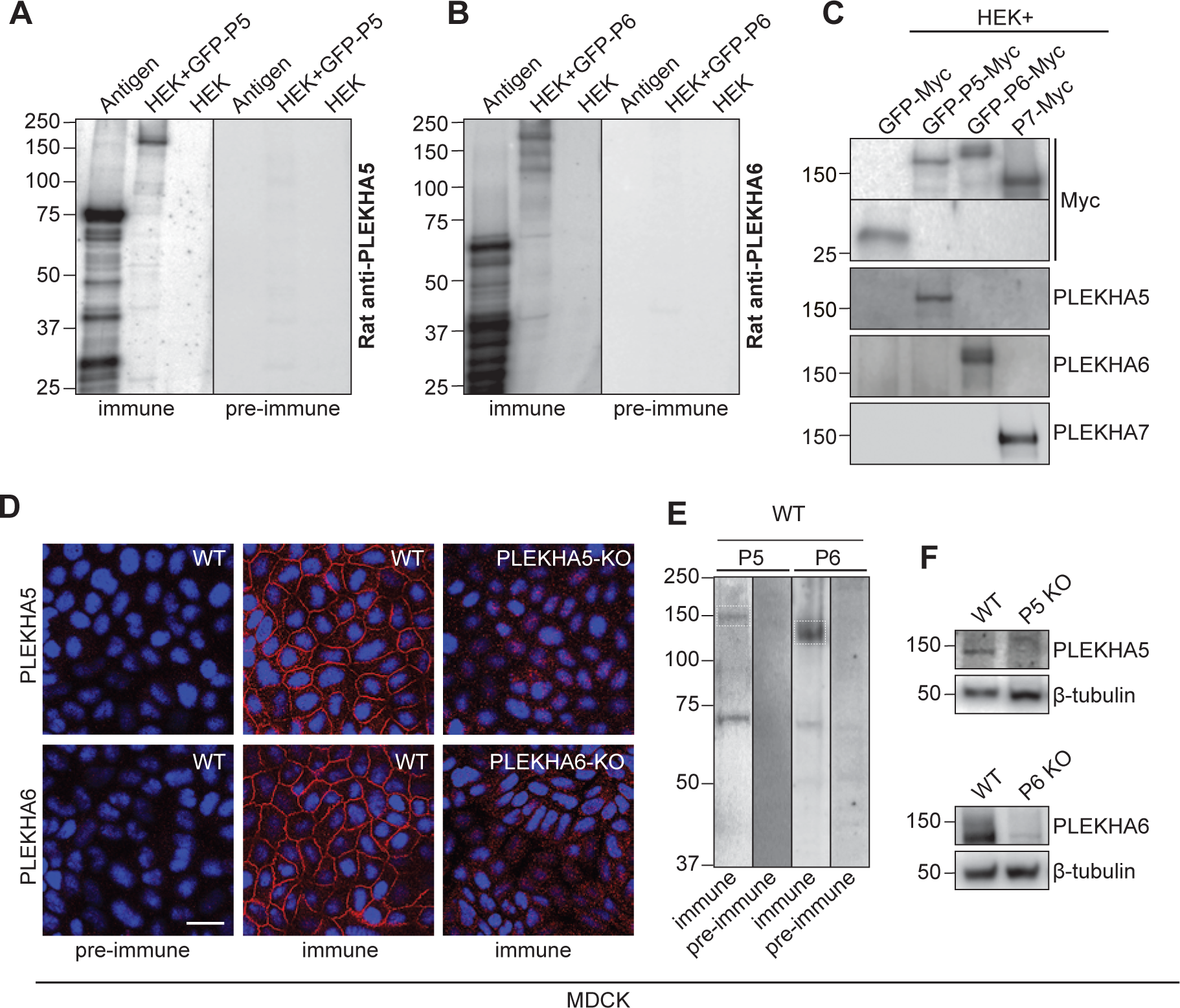
Generation and validation of antibodies against PLEKHA5 and PLEKHA6. (A) IB analysis, using anti-PLEKHA5 (left) or anti-PLEKHA6 (right) immune and pre-immune sera, of the respective antigen and of HEK cell lysates expressing the corresponding full-length protein (untransfected HEK lysate as negative control). (B) IB analysis of HEK lysates overexpressing GFP- and Myc-tagged PLEKHA5 (P5), PLEKHA6 (P6) or PLEKHA7 (P7) (GFP-Myc as control), using anti-PLEKHA5, - PLEKHA6 and -PLEKHA7 (-Myc as loading control) antibodies, showing the absence of cross-reaction. (C) IF and IB analysis of WT and KO MDCK cells (see Figure S4 for KO lines), using anti-PLEKHA5 (P5) or anti-PLEKHA6 (P6) immune and pre-immune sera. Bar= 20 µm. ß-tubulin serves as loading control.

**Figure S3.**
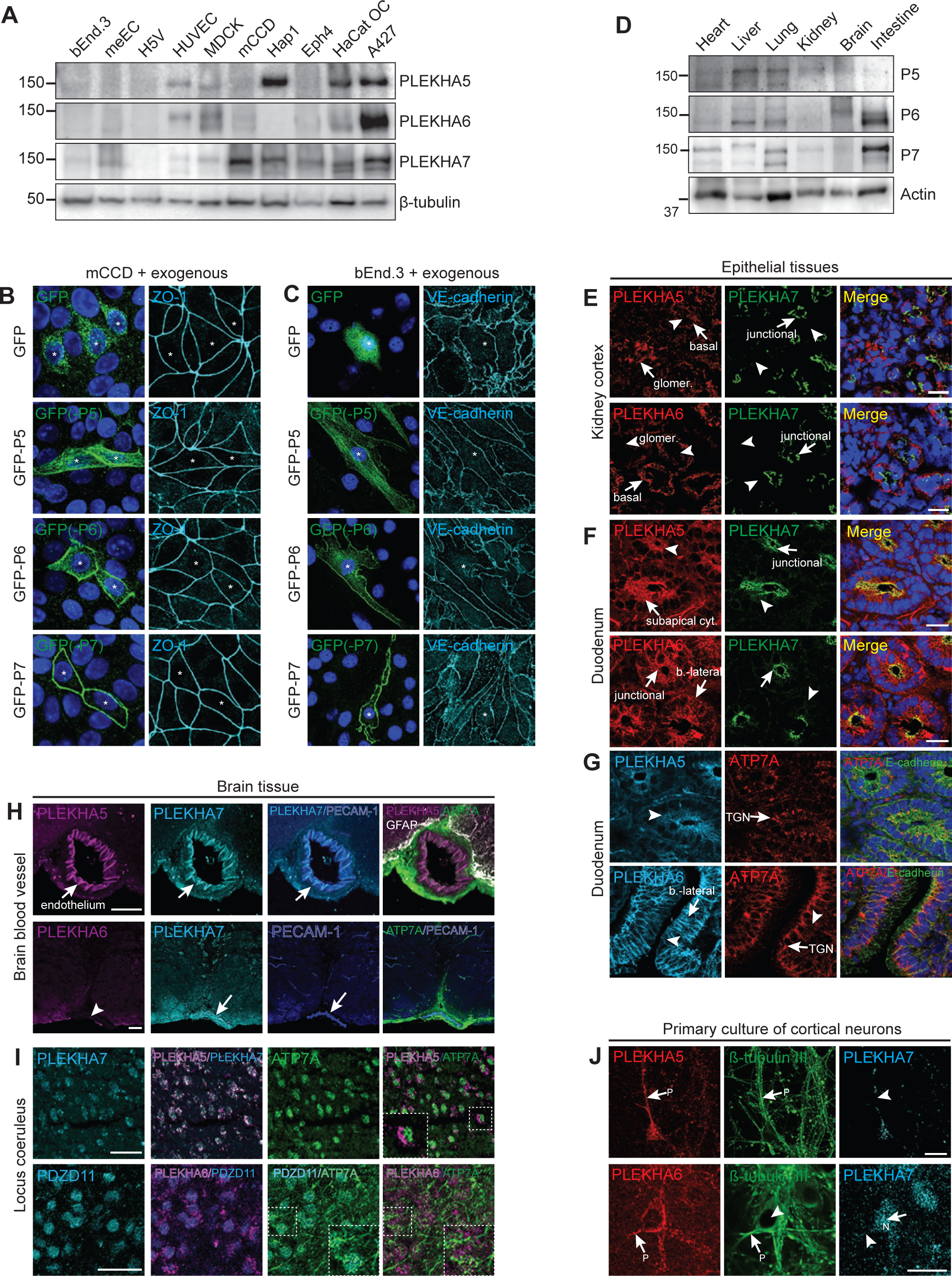
Expression and localization of PLEKHA5, PLEKHA6 and PLEKHA7 in tissues and cells. (A, D) IB analysis of PLEKHA5, PLEKHA6 and PLEKHA7 (with either ß-tubulin or actin as loading controls) in lysates of the indicated cell types (A) and mouse tissues (D). (B, C) IF analysis of GFP-tagged exogenously expressed PLEKHA5 (P5), PLEKHA6 (P6) and PLEKHA7 (P7) (GFP as control) either in epithelial mCCD (B) or endothelial bEnd.3 cells (C). Asterisks show transfected cells. (E-G) IF analysis of the localization of WW-PLEKHAs and ATP7A (indicated in each panel) in sections of mouse kidney cortex (E) and duodenum (F, G). Basal, baso-lateral (b.- lateral), junctional, glomerular (glomer.), subapical cytoplasmic (subapical cyt.) or Trans-Golgi Network (TGN) labeling is indicated by arrows; arrowheads indicate low/undetectable labeling. Bars= 20 µm. (H-I) IF labeling of mouse brain sections focusing on blood vessels (PECAM-1 as endothelial marker (blood vessels)) (H) or locus coeruleus (I). All WW-PLEKHAs and PDZD11 are expressed in neurons in locus coeruleus region, and ATP7A expression in locus coeruleus neurons appears as puncta. Bars= 50µm. (J) IF analysis of WW-PLEKHAs in primary cultures of cortical neurons, co-labelled with anti-ß-tubulin III to identify neuronal projections (pointed with P). N shows nucleus, arrows indicate labeling, arrowheads indicate low/undetectable labeling. Bars= 20 µm.

**Figure S4.**
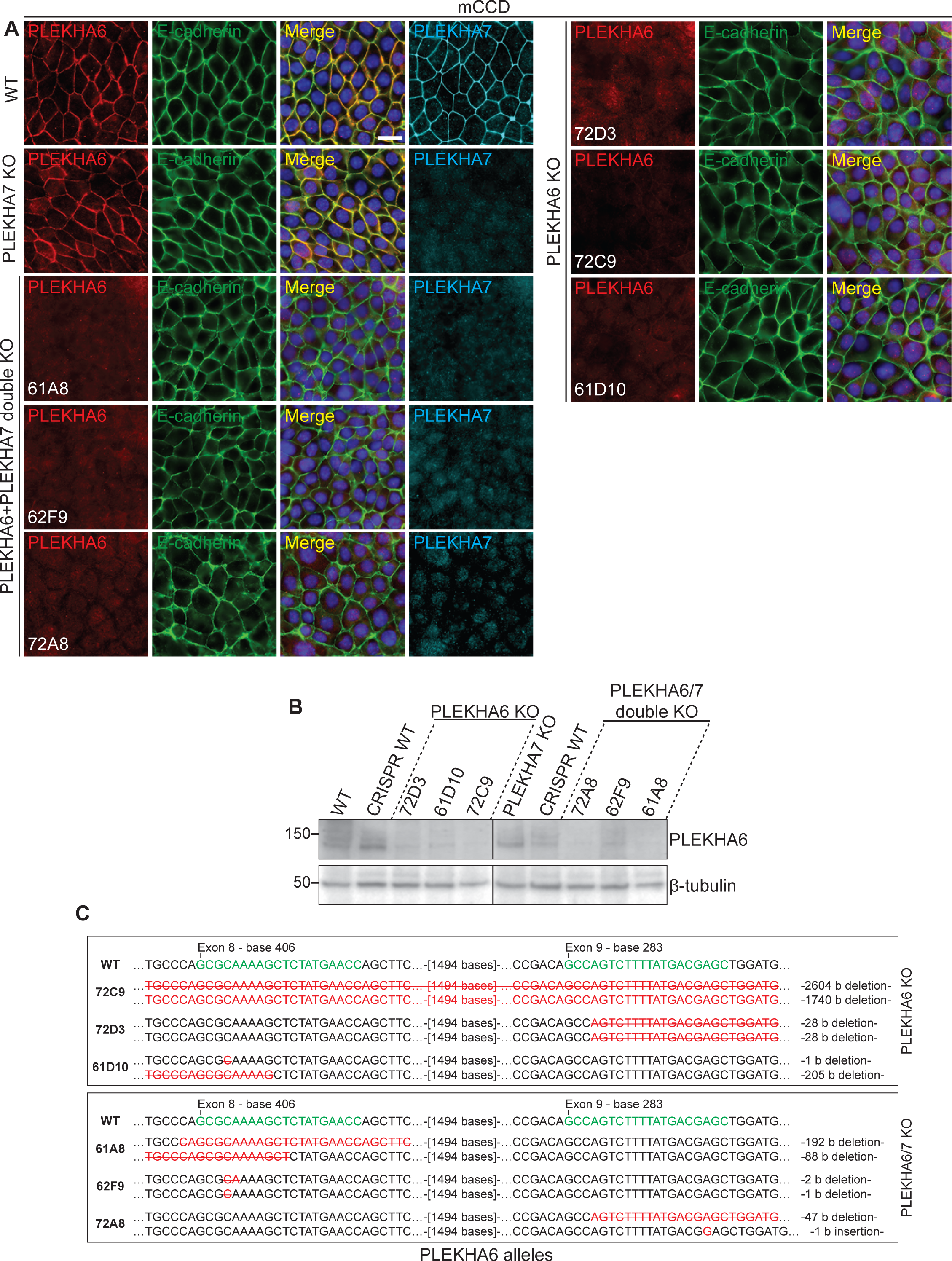
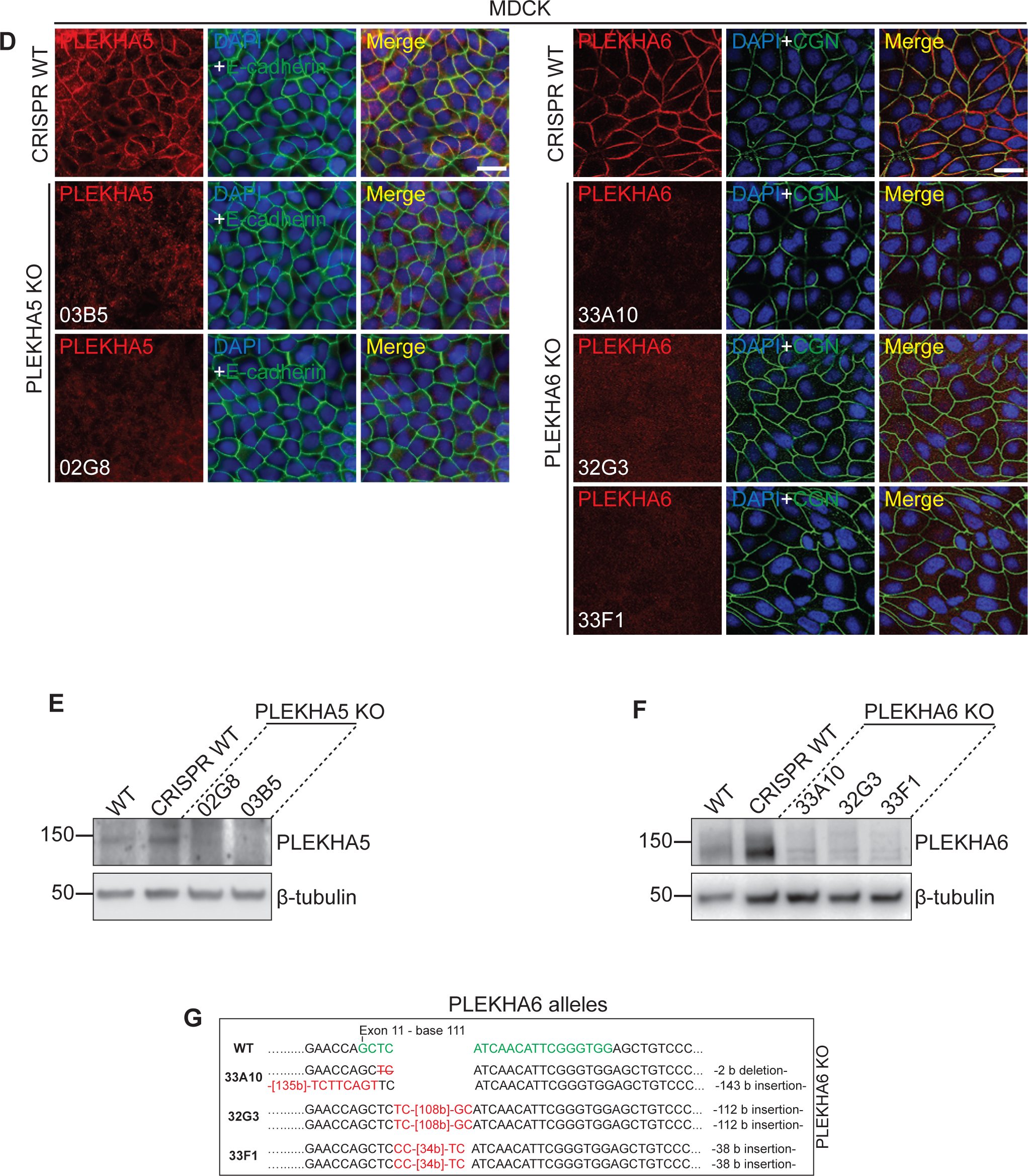
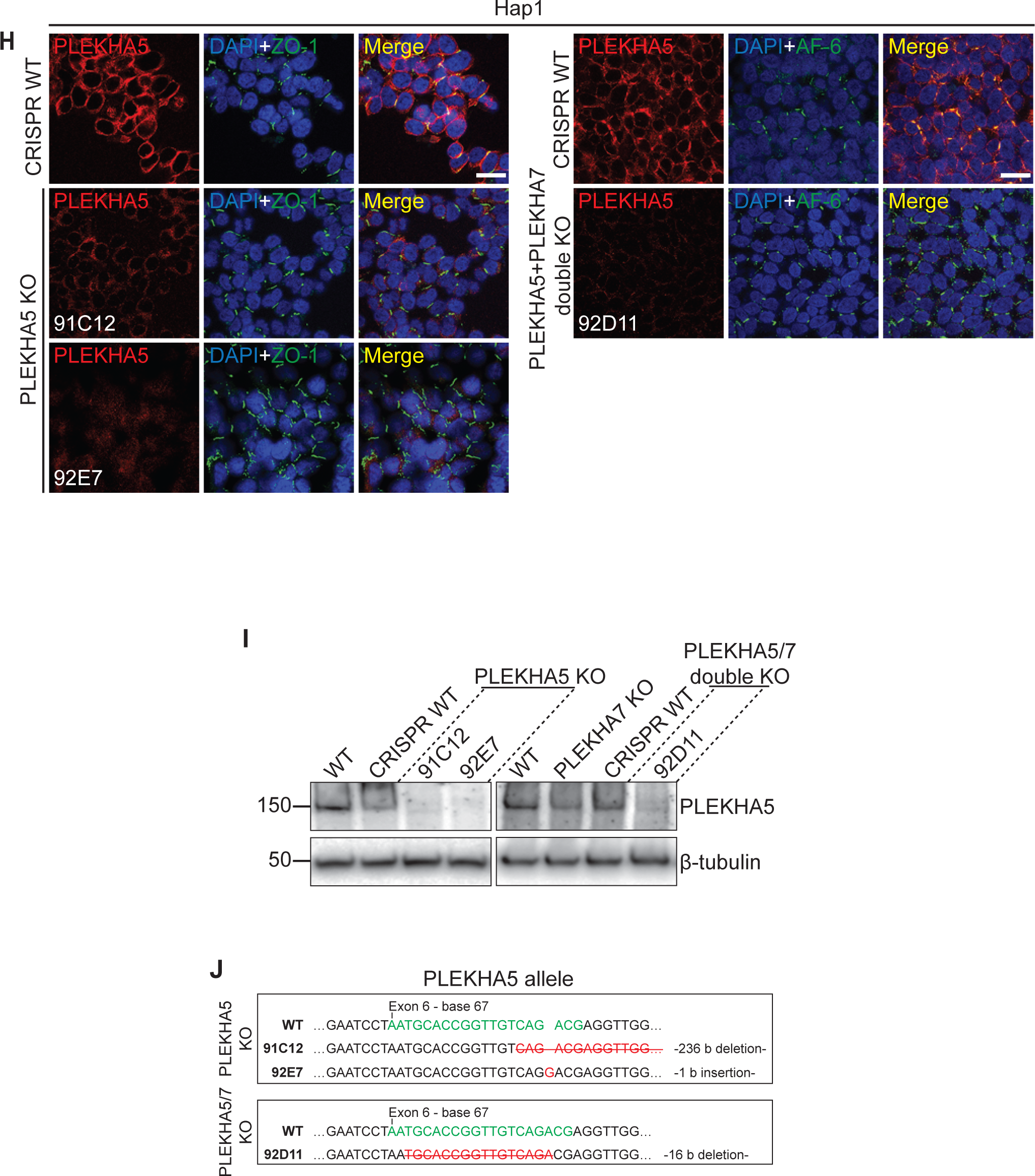
Generation of single and double PLEKHA5, PLEKHA6 and PLEKHA7 knock-out cell lines. (A-C) Validation of CRISPR/Cas9-mediated deletion of PLEKHA6 in mCCD (WT or PLEKHA7 KO background) by IF (A) and IB (B) analysis, and by genomic sequencing (C). (D-H) Validation of CRISPR/Cas9-mediated deletion of either PLEKHA5 or PLEKHA6 in MDCK by IF (D, F) and IB (E, G) analysis and genotyping (H). Since full genomic sequence for dog PLEKHA5 is not available alleles for MDCK PLEKHA5 KO clones could not be genotyped. (F-H) (I-K) Validation of CRISPR/Cas9-mediated deletion of PLEKHA5 in Hap1 (WT or PLEKHA7 KO background) by IF (I) and IB (J) analysis, and by sequencing (K). Bars= 20 µm. In C, H and K, CRISPR targets are depicted in green in the WT sequences, with their position in the exon, and respective indels in the alleles of the KO clones obtained are indicated in red.

**Figure S5.**
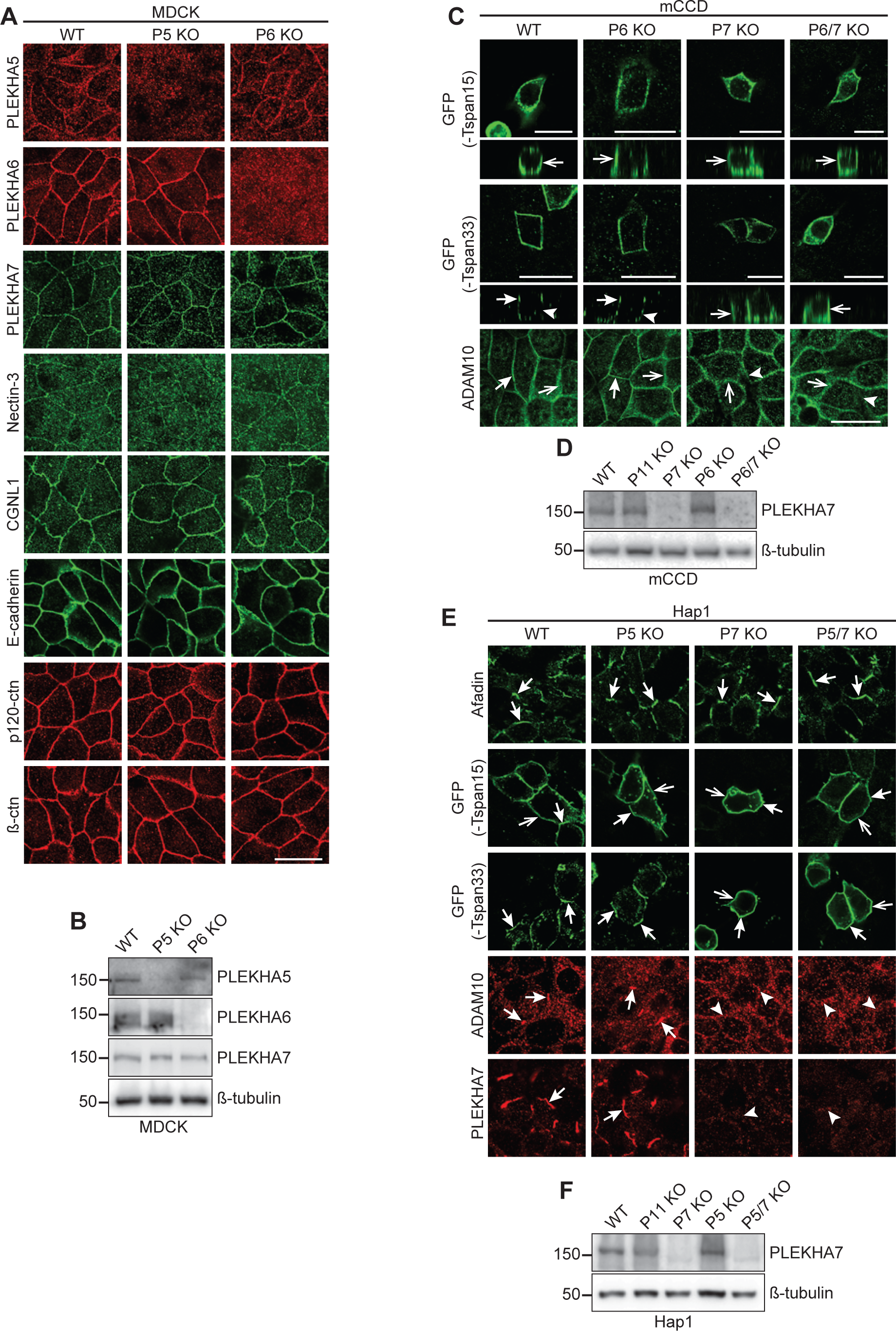
Knock-out of PLEKHA5 or PLEKHA6 does not affect the localization of cadherin complex proteins, Tspan15, Tspan33, ADAM10 and PLEKHA7. (A,C,E). IF analysis of the localizations of endogenous WW-PLEKHAs (A,E), nectin-3, paracingulin (CGNL1), E-cadherin, p120-catenin (ctn) and β−ctn (A), ADAM10 (C,E), afadin (E) and the exogenous TspanC8s Tspan33 and Tspan15 (C,E) in MDCK (A), mCCD (C) and Hap1 (E) WT and KO cells. Genotypes of KO cells are indicated on top of each column: P5=PLEKHA5, P6=PLEKHA6, P7=PLEKHA7. The phenotype of PDZD11-KO cells is identical to the phenotype of PLEKHA7-KO cells (Shah *et al*., 2018). Images showing Z section (taken at the horizontal middle position of XY view) were from cells grown on Transwells. Arrows indicate labeling, arrowheads indicate low/undetectable labeling. Bars= 20 µm. (B, D, F) IB analysis of the expression of WW- PLEKHAs in WT and KO (P5=PLEKHA5, P6=PLEKHA6, P7=PLEKHA7) MDCK (B), mCCD (D) and Hap1 (F) cells.

**Figure S6.**
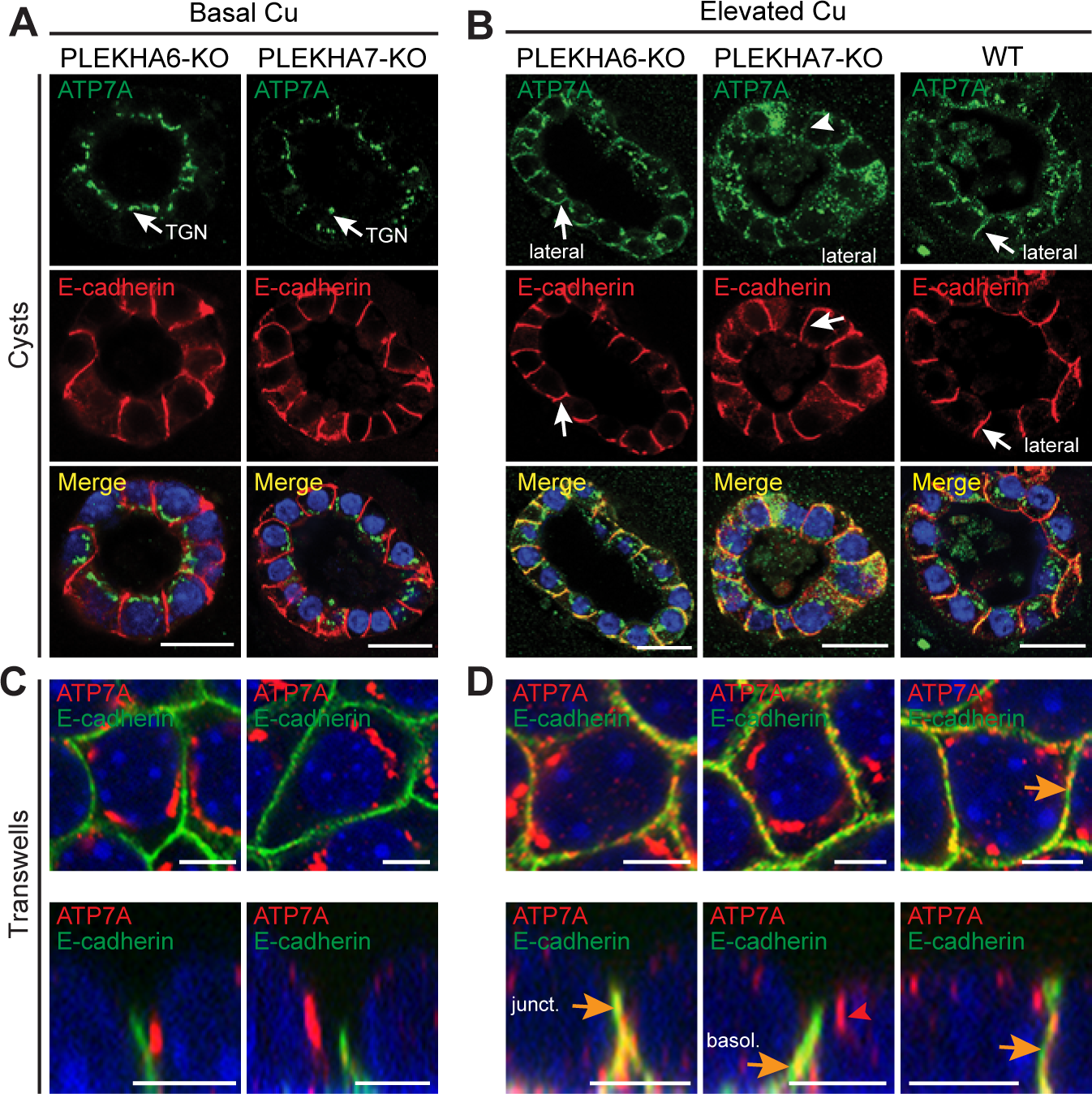
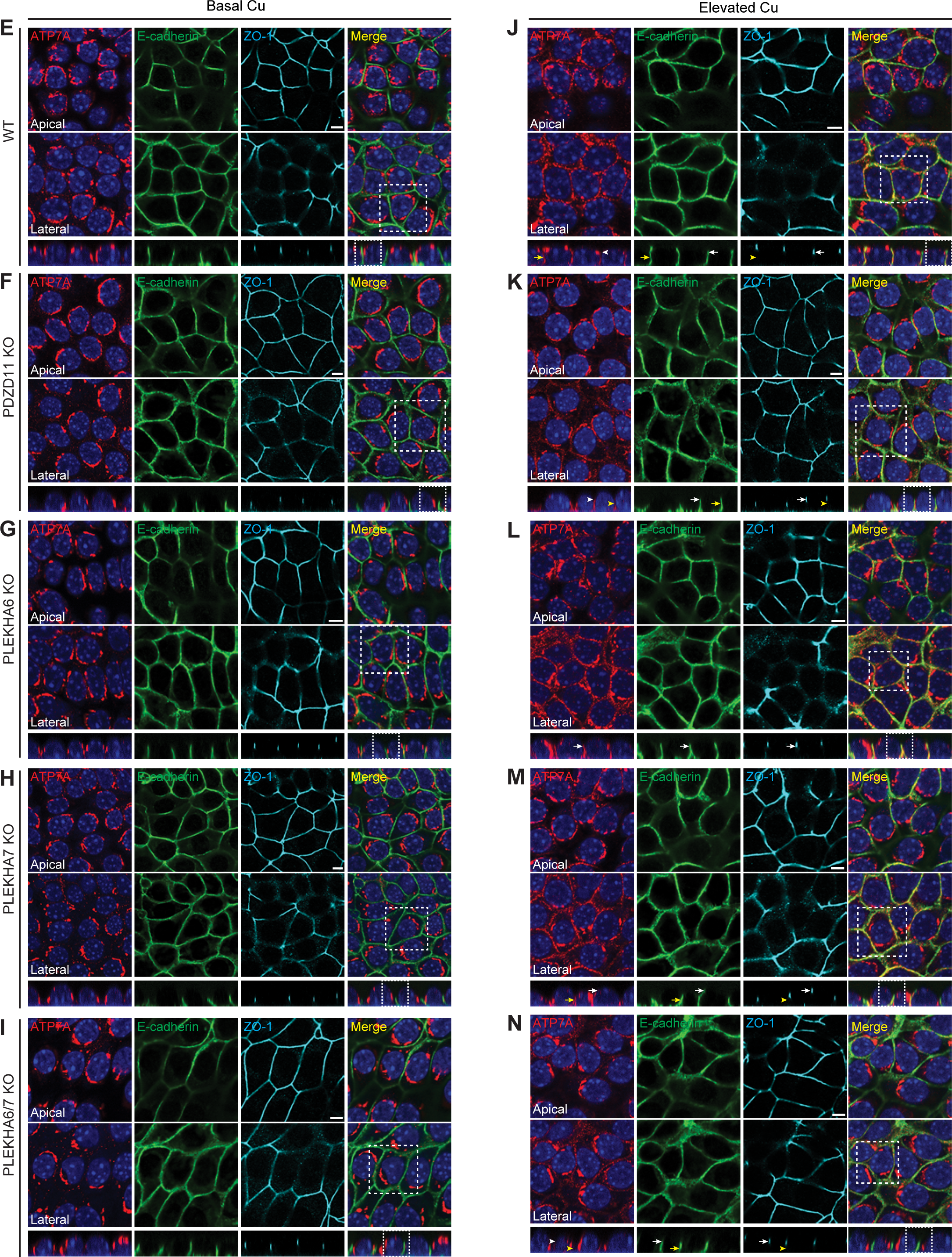
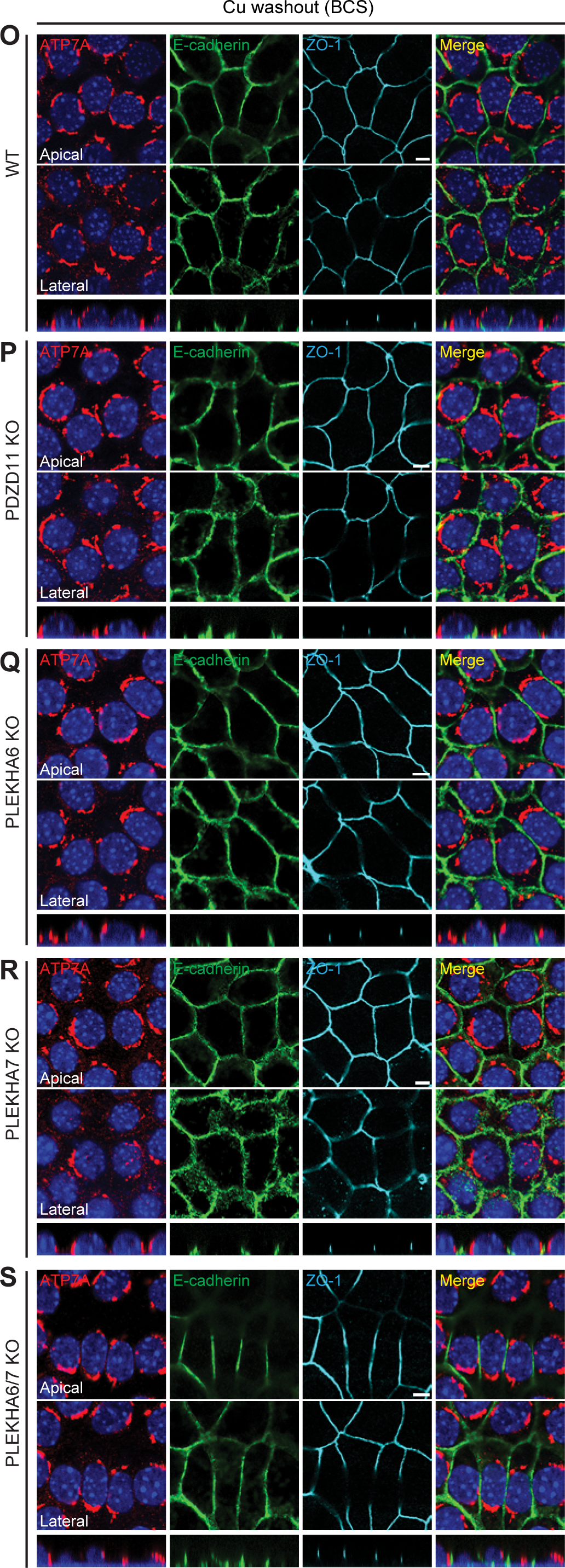
Effect of PDZD11 KO and of single and double PLEKHA6 or PLEKHA7 KO on the localization of ATP7A in mCCD cells. (A-D) IF analysis of the localization of ATP7A in PLEKHA6-KO and PLEKHA7-KO mCCD cysts (A-B) and monolayers polarized on Transwells (C-D) under basal copper conditions (A, C) or in elevated copper (B, D). TGN= Trans-Golgi Network. Lateral, apical and basal ATP7A labeling in elevated copper are indicated by arrows (B). Arrowheads indicate low/undetectable labeling. Orange arrows and red arrowheads in D indicate ATP7A labeling colocalized and non-colocalized with E-cadherin, respectively. Bars= 20 µm (A-B), 5 µm (C-D). (E-S) IF analysis of the localization of ATP7A in polarized monolayers of mCCD cells (E, J, O=WT; F,K,P=PDZD11-KO; G,L,Q=PLEKHA6-KO; H,M,R=PLEKHA7-KO; I,N,S=PLEKHA6-PLEKHA7 double KO) grown on Transwells under basal (E-I) or elevated (J-N) copper conditions, or treated with CuCl2 before copper washout (chelation with BCS) (O-S). For XY analysis, a more apical and a more basal plane of focus were imaged, using ZO-1 and E-cadherin as markers for apical junctions and lateral contacts, respectively. Merge images show colocalization between E-cadherin and ATP7A. Dotted white squares/rectangles indicate high magnification areas shown in Figure 3 and in Figure S6C-D. ATP7A labeling is detected in the TGN in all cells in basal Cu conditions and is targeted to different degrees to the cell periphery in KO cells. Copper washout resulted in the return of ATP7A to the Golgi in all cells. Bars= 5 µm.

**Figure S7.**
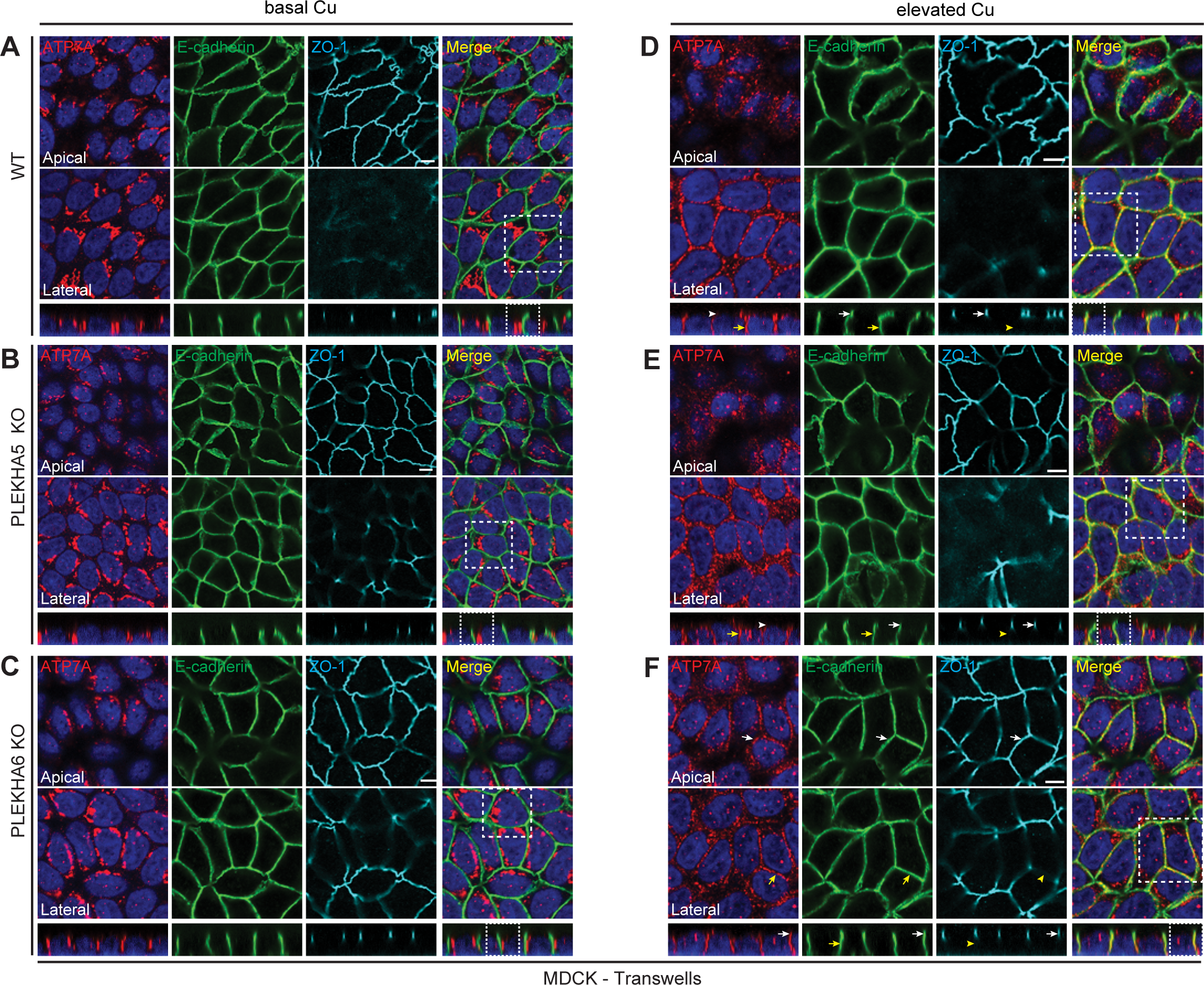
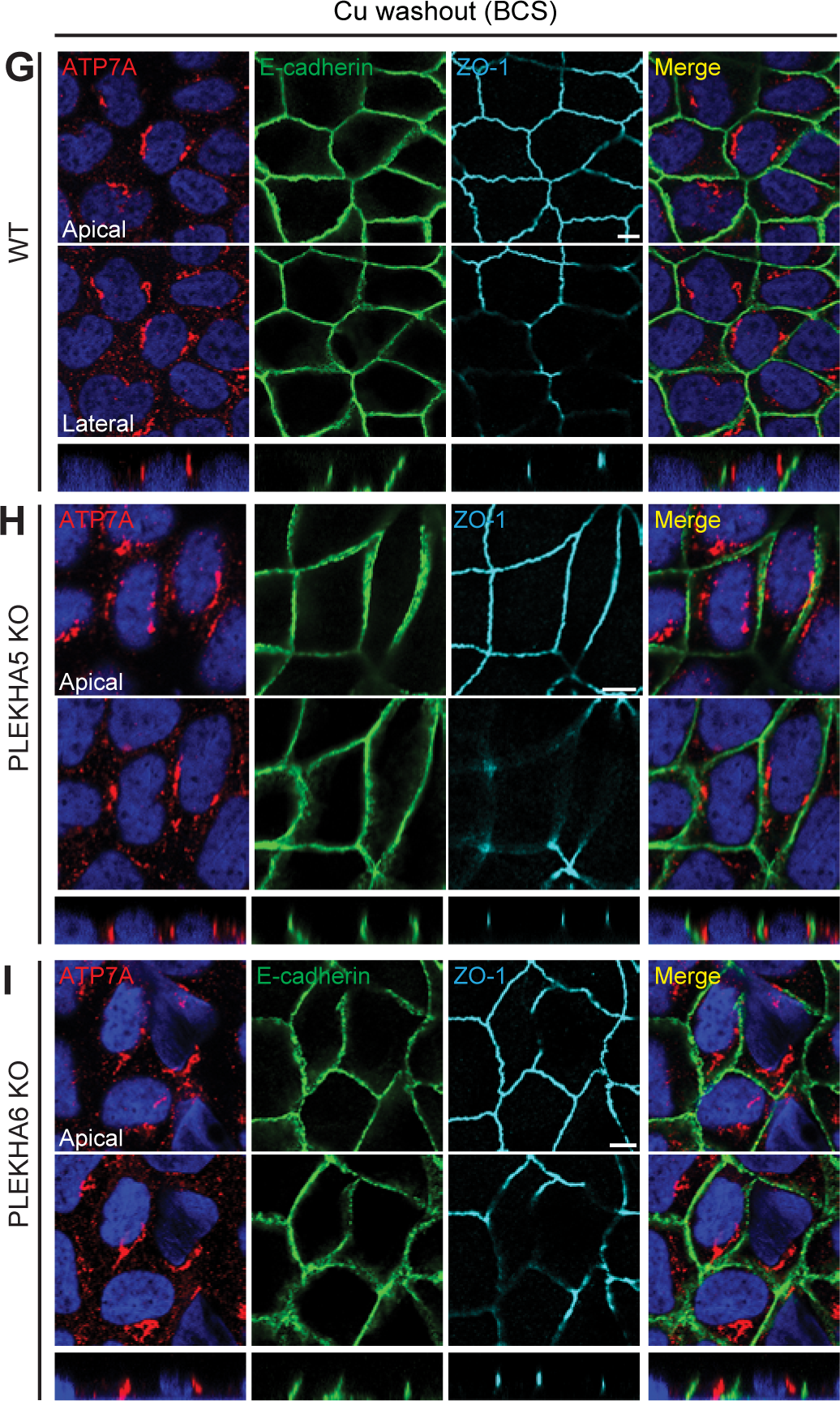
Effect of KO of either PLEKHA5 or PLEKHA6 on the localization of ATP7A in polarized MDCK cells. (A-I) IF analysis of the localization of ATP7A in polarized monolayers of MDCK cells (A, D, G=WT; B,E,H=PLEKHA5-KO; C,F,I=PLEKHA6-KO) grown on Transwells under basal (A-C) or elevated (D-F) copper conditions, or treated with CuCl2 before copper washout (chelation with BCS) (G-I). For XY analysis, a more apical and a more basal plane of focus were imaged, using ZO-1 and E-cadherin as markers for apical junctions and lateral contacts, respectively. Merge images show colocalization between E-cadherin and ATP7A. Dotted white squares/rectangles indicate high magnification areas shown in Figure 4. ATP7A labeling is detected in the TGN in all cells in basal Cu conditions and is targeted to different degrees to the cell periphery in cells KO for either PLEKHA5 or PLEKHA6. Copper washout resulted in the return of ATP7A to the Golgi in all cells. Bars= 5 µm.

